# Opposing steroid signals modulate protein homeostasis through deep changes in fat metabolism in *Caenorhabditis elegans*

**DOI:** 10.1101/551580

**Authors:** AP Gómez-Escribano, C Mora-Martínez, M Roca, DS Walker, J Panadero, MD Sequedo, R Saini, HJ Knölker, J Blanca, J Burguera, A Lahoz, J Cañizares, JM Millán, N Burton, WR Schafer, RP Vázquez-Manrique

## Abstract

Protein homeostasis is crucial for viability of all organisms, and mutations that enhance protein aggregation cause different human pathologies, including polyglutamine (polyQ) diseases, such as some spinocerebellar ataxias or Huntington disease. Here, we report that neuronal Stomatin-like protein UNC-1 protects against aggregation of prone-to-aggregate proteins, like polyQs, α-synuclein and β-amyloid, in *C. elegans*. UNC-1, in IL2 neurons, antagonizes the function of the cytosolic sulfotransferase SSU-1 in neurohormonal signalling from ASJ neurons. The target of this hormone is the nuclear hormone receptor NHR-1, which acts cell-autonomously to protect from aggregation in muscles. A second nuclear hormone receptor, DAF-12, functions oppositely to NHR-1 to maintain protein homeostasis. Transcriptomics analyses reveal deep changes in the expression of genes involved in fat metabolism, in *unc-1* mutants, which are regulated by NHR-1. This suggest that fat metabolism changes, controlled by neurohormonal signalling, contributes to modulate protein homeostasis.

## BACKGROUND

Protein homeostasis is essential to maintain appropriate cell function. This consists of both, an adequate expression of specific sets of proteins and proper folding and localisation of these molecules. When proteins do not fold correctly they tend to present to the cytoplasm hydrophobic domains, which normally lay in the interior of the protein are exposed to the cytoplasm, and hence become prone to aggregation. These aggregation processes produce very toxic species that substantially alter the function of cells and cause many neurodegenerative diseases such as Huntington disease (HD), together with eight polyglutamine-associated disorders, Alzheimer’s, Parkinson’s and ALS ^1^. To prevent the formation of aggregates there are natural mechanisms to control proper folding of proteins and to remove proteins that are already misfolded ^2^. Once a considerable amount of proteins has collapsed, due to improper folding, they induce stress in the cytoplasm, by poisoning the machinery of macroautophagy (autophagy) or the proteasome system. This in turn activates unfolded protein response (UPRs) pathways in the endoplasmic reticulum, mitochondria and cytosol ^3^. UPR signals activate nuclear transcription factors that induce expression of genes associated with folding and degradation of proteins, autophagy, among other protective pathways ^3^. In addition, chemical synaptic function in neurons was shown to regulate polyglutamine (polyQ) misfolding in muscle cells ^4,5^. In this regard, Silva and co-workers showed that a small increase in physiological cholinergic signalling, at the motor synapses, induced a calcium-dependent activation of muscular HSF-1, which in turn activates the expression of chaperones that protect against protein aggregation ^5^. Despite these and similar observations, the mechanisms by which neuronal signalling can control protein aggregation in distant tissues, remains unknown.

Nuclear receptors comprise a group of transcription factors that regulate the expression of genes in a ligand binding-dependent manner ^6,7^. These ligands are usually lipid molecules, such as steroid hormones, vitamins (D_3_ and A), metabolites and xenobiotics ^8^. In some cases, these molecules need to be processed by sulfotransferases, which add a sulphate moiety, and sulfatases, which desulfate the molecule allowing it to bind the receptor, to promote hormonal signalling to distal tissues. *C. elegans* genome encodes one sulfotransferase, SSU-1, which is expressed exclusively in ASJ neurons ^9^, and three sulfatases, SUL-1, SUL-2 and SUL-3 ^10^. The *C. elegans* genome encodes 284 NR genes in contrast with humans which have only 48 ^11^. One of the best-known nuclear receptor in *C. elegans*, DAF-12, regulates lipid metabolism, lifespan and development ^12–14^ through the dafachronic acid hormones ^15^. This control depends on the tight modulation of the expression of genes encoding key metabolic enzymes, which includes negative feedback loops operated by their enzymatic products ^16^. For example, NHR-49, a homologue of the Hepatocyte Nuclear Factor 4-alpha (HNF4α) ^17^, has a crucial role to regulate lipid metabolism, longevity and nutrient response ^18,19^. After activation, nuclear receptors dimerise, producing homo- and heterodimers, and bind DNA domains in the vicinity of promoters to regulate gene expression, sometimes in opposite manner, depending of the NR partner. For instance, heterodimeric NHR-49/NHR-80 promotes lipid desaturation while NHR-49/NHR-66 blocks sphingolipid metabolism and lipid remodelling ^20^. Among the lipids regulated by these NRs, oleic acid has been shown to promote longevity in germline ablated mutants ^21^. This lipid also is able to enhance proteostasis, in worms expressing polyQs, promoted by XBP-1 signalling ^22^. Oleic acid is synthesised by several key enzymes, that are encoded in the *C. elegans* genome: FAT-6 and FAT-7 ^23^, whose expression is regulated by NHR-80 ^24^.

Here we report the isolation of a loss-of-function allele of *unc-1* in *C. elegans*. This gene encodes a homologue of the mammalian Stomatin-like protein. Depletion of *unc-1* alters motor coordination and enhances polyQ aggregation in worms. We show that UNC-1 functions in neurons, likely by controlling electrical synapse, to regulate protein homeostasis by cell-nonautonomous signalling. Disruption of *unc-1* function causes a malfunction of the electrical synapse between IL2 and ASJ neurons, which likely secretes an excess of a sulphated signal, which activates NHR-1 in muscle cells and neurons. Over-activation of NHR-1, in *unc-1(vlt10)* animals, down-regulates genes encoding enzymes of the fat metabolism, which in turns disrupts proteostasis. In contrast, we show that signalling through DAF-12 functions antagonistically to NHR-1, to control protein aggregation. These results evidence for the first time that nuclear receptors modulates polyQ aggregation through lipid metabolism remodelling. Moreover, some of the enzymes involved in the signalling pathway leading to NHR-1 activation are potential druggable targets, that may be used to treat neurodegenerative diseases caused by protein homeostasis disruption.

## RESULTS

### A forward genetic screen identifies *unc-1* as a modulator of polyQ aggregation

García and co-workers have shown that extracellular signalling modulates protein homeostasis ^4^. However, upstream and downstream processes involved in this regulation, are not completely understood, neither regarding the existence of other synaptic-related mechanisms that may influence protein homeostasis. To expand our knowledge on these signalling pathways controlling protein homeostasis we performed an EMS mutagenesis screen for genes regulating protein aggregation (Fig. 1A), in a worm model of polyQ diseases ^25^. We used animals that express 40 glutamines fused to a yellow fluorescent protein (*40Q::YFP*) in muscle cells ^25^. In these animals, 40Q aggregates, in an age-dependent manner, and forms inclusion bodies, that can be observed and counted under a dissecting microscope. Among several mutants, we obtained a mutant strain, RVM10 that carries the allele *vlt10*, which substantially enhanced the aggregation pattern (Fig. 1B, C) without affecting *40Q* transgene expression (Supplementary Fig. 1A, B). In addition to enhanced polyQ aggregation, the *vlt10* animals showed an uncoordinated phenotype, suggesting that this mutation may affect the synaptic function of the nervous system and/or the muscle cells.

**Figure 1.**
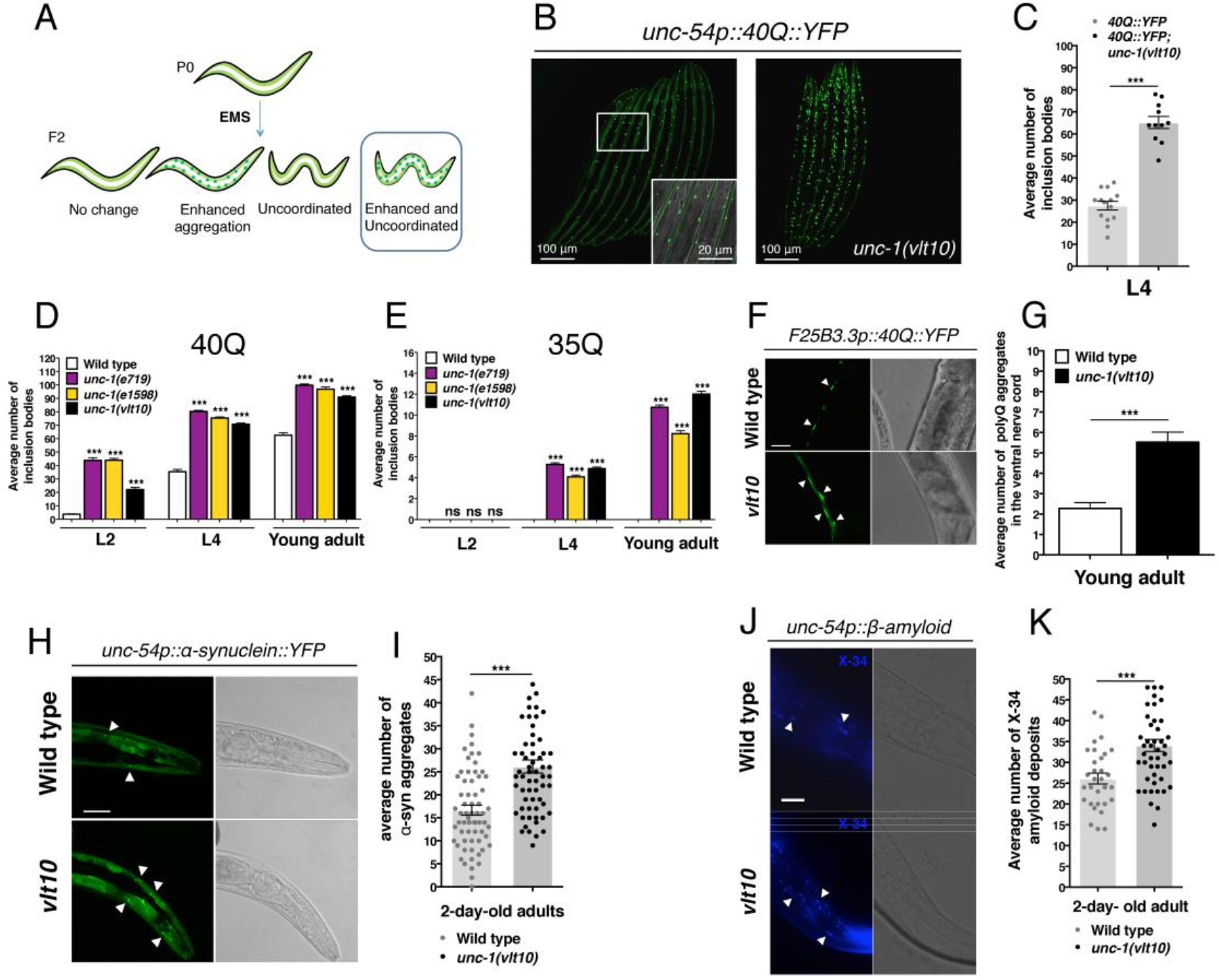
UNC-1 modulates protein aggregation. (**A**) Isolation of a mutant strain, which showed uncoordination and altered polyQ aggregation, after treatment with ethyl methanesulfonate *40Q* animals. (**B**) Representative images of L4 wild type and *unc-1* mutant worms, obtained by confocal microscopy. The white box shows an area that has been further magnified 20 times, to show the shape and pattern of the inclusion bodies in muscle cells (white arrowheads). (**C**) Animals that have the homozygous *vlt10* allele show increased polyQ aggregation. (**D**) The *e719* and *e1598* alleles of *unc-1* emulate aggregation of *vlt10*, at several stages. (**E**) *vlt10, e719* and *e1598* alleles enhance polyQ aggregation in *35Q* animals. (**F–G**) *unc-1* regulates proteostasis in the nervous system. Representative images from fluorescence microscopy showing neuronal aggregates of polyQs (white arrowheads) in the ventral nerve cord of control *F25B3*.*3p::40Q::YFP* and mutant animals *F25B3*.*3p::40Q::YFP*; *unc-1 (vlt10)* in the young adult stage. *vlt10* increases the number of neuronal aggregates of polyQs. (**H–I**) *unc-1(vlt10)* enhances aggregation of α-synuclein. Representative images from confocal microscopy showing the area of the muscle tissue located between the two pharyngeal bulbs, where the measure of α-synuclein aggregates has been obtained. White arrowheads indicate α-syn aggregates. (**J–K**) Depletion of *unc-1* enhances β-amyloid deposits in two-day-old adults. Representative images taken with fluorescence microscopy that include the area between the two pharyngeal bulbs, where are highlighted amyloid deposits (white arrows). The plotted data show the mean ± SEM. The tested at least 30 animals for each specific stage for all graphs, except for graph G (60 animals). Each analysis has been reproduced at least three times. ***: p-value < 0.001; ns: statistically not significant. Statistical t-test with non-parametric Mann-Whitney test (graph C, G, I and K). Statistical test ANOVA with multiple comparative test of Tukey (graph D-E). Scale bar: 20 μm.

To determine the molecular identity of this allele we sequenced the whole genome of this strain after six outcrossing steps and also the genome of the original strain (RVM10). EMS causes random lesions through the chromosomes and after outcrossing RVM10, most of these mutations would be lost during outcrossing, except the DNA changes that lie around the allele responsible for the aggregation phenotype. We took advantage of this, and when we compared both genomes we observed a region with dense amount of mutations that “peaked” around the left arm of chromosome X (Supplementary Fig. 1C). Detailed analysis of this region showed that the *unc-1* gene, of RVM10, had a nonsense mutation in homozygosis that gives rise to premature stop codon, which would produce a putative null (Supplementary Fig. 1D). *unc-1* encodes a homologue of the Stomatin-like protein family from mammals ^26,27^. The product of this gene has been shown to be involved in modulation of the electrical synapse ^28^ and sensitivity to anaesthetics ^27^. Null alleles of this gene are well-known to cause strong uncoordination ^28^, similar to *vlt10* mutant from our screen.

### Depletion of *unc-1* enhances polyQ aggregation

To test whether the loss of function of *unc-1* is responsible for the enhanced aggregation phenotype of *vlt10* mutants, we introduced well-known mutant alleles of *unc-1* (*e719* and *e1598*) ^27,29^ into the *35Q* and *40Q* worms. In both cases, analysis of the inclusion bodies produced by the aggregation of polyQs showed that mutations in *unc-1* (*e719* and *e1598*) phenocopy the aggregation pattern of *vlt10* worms (Fig. 1D, E). In addition, we investigated whether *unc-1(vlt10)* would modify the late aggregation phenotype of a strain expressing *35Q* fused to YFP (*35Q::YFP*) in body wall muscles. These worms have a later and weaker phenotype than animals expressing *40Q*, and they produce inclusion bodies at late adult stages ^25^. Analysis of *35Q*; *unc-1(vlt10)* animals showed that the *vlt10* allele is able to enhance and accelerate the phenotype of these animals to a similar extent as other *unc-1* alleles, *e179* and *e1598* (Fig. 1E). These results indicate that *unc-1* is a modulator of polyQ aggregation in muscle cells.

As *unc-1* is expressed also in neurons, we sought to investigate whether *vlt10* may affect polyQ aggregation in this tissue. To do so we used a worm strain that expresses *40Q* in the whole nervous system ^30^, in which we introduced the mutant allele *vlt10*. As we expected, *unc-1(vlt10)* increased neuronal aggregates formation in the ventral nerve cord of young adults, further demonstrating the requirement of *unc-1* in protein homeostasis maintenance (Fig. 1F, G). Altogether, these data indicate that *unc-1* is required to prevent aggregation of polyQ-containing proteins both in muscle cells and neurons.

### *unc-1(vlt10)* enhances α-synuclein and β-amyloid aggregation

Having shown that *unc-1* regulates polyQ aggregation we sought to test whether its function is polyQ-specific or it may affect other prone-to-aggregation proteins, such as α-synuclein and β-amyloid. These proteins aggregate in brains of patients of Parkinson and Alzheimer diseases, respectively ^31^. To do so we introduced the *unc-1(vlt10)* allele into transgenic worms expressing α-synuclein::YFP (from now on α-syn) ^32^ and worms expressing human β-amyloid ^33^, in body wall muscles. In contrast with worms expressing polyQs, which show aggregation early during the life cycle of worms, both α-syn and β-amyloid aggregates appear during late stages of adulthood (2-day-old adults). Analysis of these worms showed that *unc-1* loss of function increased α-syn and β-amyloid aggregation in 2-day-old adults compared to controls (Fig. 1H-K). These data suggest that *unc-1*, rather than being polyQ-specific, is a general modulator of protein homeostasis.

### *unc-1* is required in the nervous system to maintain protein homeostasis non-cell-autonomously

UNC-1 is expressed both thorough the nervous system and in muscles cells ^28,34,35^. To investigate whether UNC-1 modulates muscle aggregation in a cell-autonomous manner, we reintroduced the cDNA of *unc-1*, under the control of the promoter of *myo-3* (*myo-3p*), a gene that is exclusively expressed in muscle cells ^36^ in *40Q*; *unc-1(vlt10)* animals. Analysis of these worms showed that this construct seemed to partially rescue aggregation of polyQs (Supplementary Fig. 2A). However, we noticed that these transgenic animals showed an obvious reduction of YFP expression, to the naked eye. We hypothesised that the presence of high copy numbers of the *myo-3* promoter may be interfering with the expression of the *40Q* insertion (Supplementary Fig. 2B). In support of this hypothesis, we observed a reduction of inclusion bodies when we injected the *myo-3* promoter alone (data not shown). Real time PCR experiments on the YFP gene further demonstrated that these worms had a clear reduction of *40Q* expression (Supplementary Fig. 2B). Therefore, it is likely that the apparent rescue observed is just an artefact due to a reduction of transgene expression, and not to a restoration of *unc-1* function in muscle cells. However, we cannot rule out that at least part of the effect is genuine rescue.

When we expressed *unc-1* in the nervous system, using the pan-neuronal promoter of the *rab-3* gene ^37^, *unc-1(vlt10)* animals showed a substantial reduction of polyQ aggregation in all stages (Fig. 2A). In contrast with the attempt of rescuing *unc-1* deficiency in muscle cells, specific expression of *unc-1* in the nervous system did not modify the expression of *40Q* (Fig. 2A). To confirm these results, we knocked-down *unc-1* expression using RNAi in a tissue-specific manner using transgenesis ^38,39^ (Methods and Supplementary Fig. 8). We used RNAi against the ampicillin resistance gene (*AMP*^*r*^) as a negative control. Tissue-specific silencing of *unc-1* showed an increase of the number of inclusion bodies compared with *40Q* and *40Q*; *AMP*^*r*^*(RNAi)* animals (Fig. 2B), further suggesting that aggregation of polyQ in muscle cells is controlled from neural *unc-1* expression. Finally, to undoubtedly show that the function of *unc-1* is purely neuronal, we used a null dominant allele of *unc-1, n494*, to disrupt *unc-1* function within the nervous system. Analysis of worms expressing a cDNA of *unc-1* carrying this allele showed that these animals had a similar level of inclusion bodies than *40Q*; *unc-1(vlt10)* animals (Fig. 2C). Moreover, we used *unc-1* overexpression to show that it is not an overload of UNC-1 which induces an increase of polyQ aggregation (Fig. 2C). Altogether these results show that UNC-1 modulates polyQ aggregation in muscle cells in a cell-nonautonomous fashion from neurons, probably by modulating the gap junctions (Table 1 and Supplementary Fig. 3). In support of these data, we also analysed the aggregation patterns of worms expressing muscular *40Q*, in which we depleted different gap junction components (innexins) and other stomatin like proteins. These experiments showed that alteration of some of these components indeed increase polyQ aggregation in muscle cells (For more information to see Supplementary Fig. 3).

**Figure 2.**
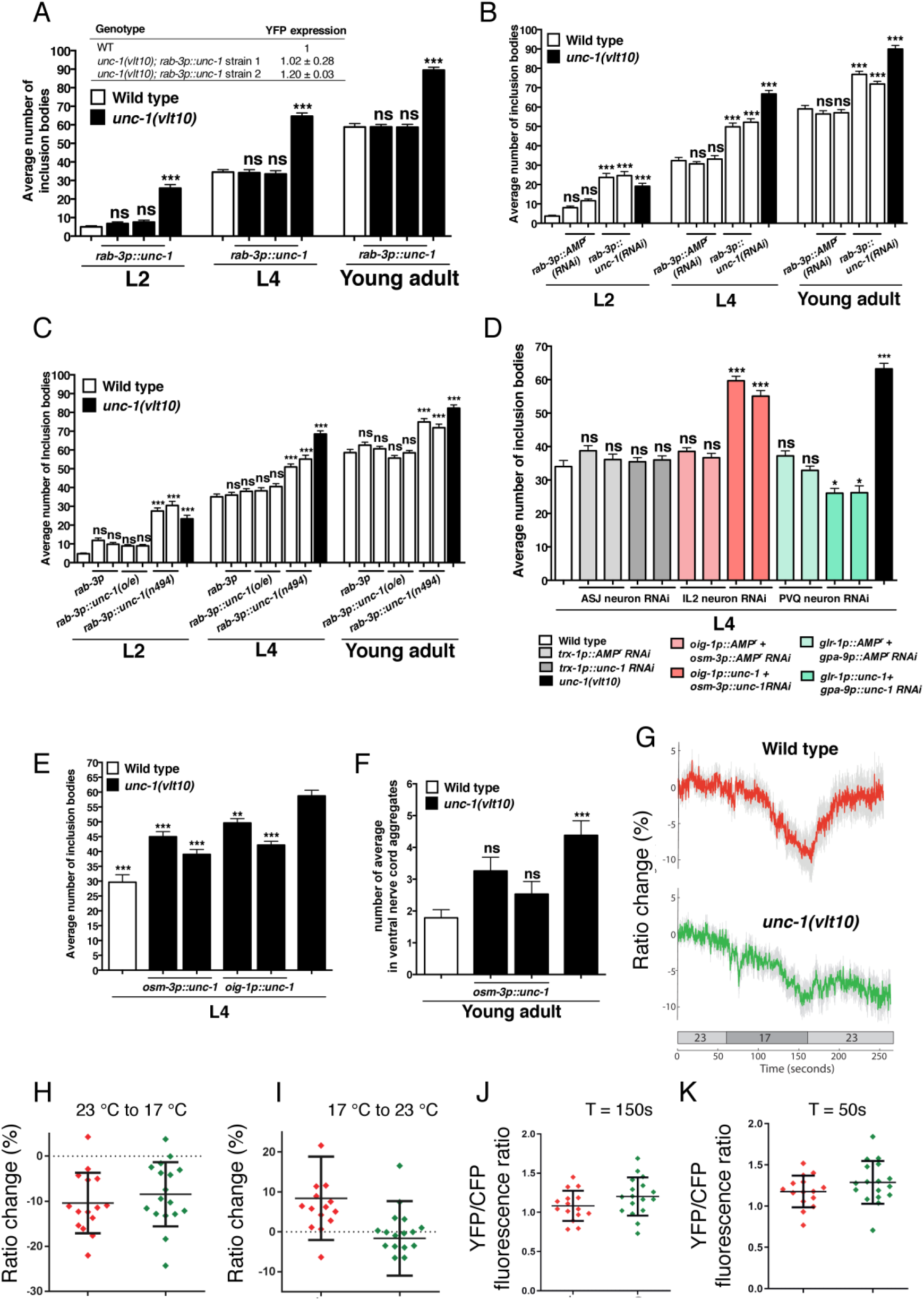
UNC-1 is required in IL2 neurons to modulate protein homeostasis in muscle cells and neurons. (**A**) Expression of the cDNA of *unc-1* in the whole nervous system rescue polyQ aggregation in *unc-1* mutants, at several stages (L2, L4 and young adult). Tissue-specific rescue of *unc-1* does not affect the expression of the *40Q:: YFP* transgene (table inserted in the graph). (**B**) Tissue-specific RNAi against *unc-1* in the nervous system increases the average number of polyQ inclusion bodies in muscle cells. (**C**) Overexpression of *unc-1* in the nervous system has no effect on aggregation of polyQs. The expression of the dominant negative allele of *unc-1, n494*, in the nervous system, increases polyQ aggregation. (**D**) specific RNAi against of *unc-1* in IL2 neurons, emulates pan-neuronally-induced RNAi against *unc-1*, while specific silencing of *unc-1* in ASJ does not produce changes in aggregation of polyQs. PVQ neuron-specific *unc-1* silencing reduces weakly polyQ aggregation compared to controls. (**E-F**) Restoration of the *unc-1(cDNA)* in *unc-1(vlt10)* mutants using either *osm-3p* or *oig-1p*, significantly rescues the *unc-1* associated-polyQ aggregation in muscle cells and neurons. (**G-K**) *unc-1(vlt10)* alters ASJ response to temperature. Calcium imaging in ASJ, in response to temperature shift. (**G**) Average traces of % YFP/CFP fluorescence ratio change. Grey indicates SEM, bar at the bottom indicates the perfusion temperature. **(H-I)** Scatter plots showing individual ratio changes in response to the temperature shift from 2 3°C to 17 °C (comparing the mean YFP/CFP fluorescence ratio at T = 50s with T = 150s, using a 20-frame window) and from 17 °C to 23 °C (comparing YFP/CFP fluorescence ratio at T = 150s with T = 150s). (**J-K**) Scatter plots showing the YFP/CFP fluorescence ratio at T = 50s with T = 150s. Bars indicate mean ± SEM. thirty animals were tested per strain and/or condition. The analysis has been reproduced at least three times. ***: p-value < 0.001; **: p-value < 0.01; *: p-value < 0.05; ns: statistically not significant. Statistical test ANOVA with multiple comparative test of Tukey.

**Table 1.** Stomatin-like proteins and innexins are required to maintain protein homeostasis.

| Genotype | Focis (%)<sup>1</sup> | P value<sup>2</sup> | Protein | Function<sup>3</sup> | Expression<sup>3</sup> |
| --- | --- | --- | --- | --- | --- |
| *unc-1(vlt10)* | 171.7<br>± 3.1 | *** | Stomatin-like protein | Locomotion<br>Sensitivity to volatile anaesthetics | NS-WE<sup>4</sup> |
| *unc-24(e138)* | 149.1<br>± 3.8 | *** | Stomatin-like protein | Locomotion<br>Sensitivity to volatile anaesthetics | NS-WE |
| *mec-2(e75)* | 114.5<br>± 5.2 | ns | Stomatin-like protein | Mechanosensation | Mechanosensory neurons |
| *unc-7(e5)* | 178.7<br>± 4.7 | *** | Innexin | Locomotion<br>Sensitivity to volatile anaesthetics<br>Sensitivity to ivermectin | NS<sup>4</sup><br>Glial cells<br>Mesoderm |
| *unc-9(ec27)* | 185.4<br>± 2.7 | *** | Innexin | Locomotion<br>Sensitivity to volatile anaesthetics<br>Sensitivity to ivermectin<br>Electrical coupling of muscle cells | NS-WE<br>Enteric muscles<br>RS<sup>4</sup><br>Hypodermis<br>Glial cells<br>Embryos |
| *inx-2(vlt22)* | 175.4<br>± 4.3 | *** | Innexin | Role initial cellular proliferation stage | NS-AVK neuron<br>Pharynx<br>Intestine<br>Embryos |
| *inx-7(ok2319)* | 98.7<br>± 3.8 | ns | Innexin | Unknown | NS-WE<br>Pharynx<br>RS<br>Mesoderm<br>Embryos |
| *inx-6(rr5)* | 98.8<br>± 4.1 | ns | Innexin | Synchronized pharyngeal muscle contractions | Pharynx<br>Mesoderm<br>Embryos |

<sup>1</sup>Average of inclusion bodies normalised with the background of wildtype siblings ±Standard Error of the Mean.<sup>2</sup>ANOVA with post-hoc analysis of Tukey: \*\*\*p<0.001 referred to wild type; ns: no statistically significant.<sup>3</sup>Function and expression according by the review of <sup>105</sup>. <sup>4</sup>NS-WE: nervous system-wide expression; NS: some neurons; RS: reproductive system.

### *unc-1* is required in IL2 neurons to modulate polyQ aggregation

Previous reports showed genetic interaction between *unc-1* and *ssu-1* regarding anaesthetics and motor phenotypes in worms ^9^. Since *ssu-1* is expressed exclusively in ASJ ^9^, it is tempting to speculate that the function of *unc-1* is required in this cell to modulate polyQ aggregation. However, when we tried to rescue *unc-1(vlt10)* by expressing *unc-1* cDNA under the control of the *trx-1* promoter, which expresses only in ASJ neurons, we did not observe any change in 40Q aggregation (data not shown). Since UNC-1 regulates electric synapses, we hypothesised that *unc-1* may be required in neurons that connect ASJ through gap junctions. In this regard, Cook and co-workers published a detailed description of the *C. elegans* connectome, where they show all nerve connexions of electrical and chemical synapses of the worm ^40^. They show that ASJ makes electrical synapsis with IL2L and PVQR ^40^. Hence, we sought to disrupt the function of *unc-1* in ASJ, IL2L and PVQR using RNAi induced from transgenes ^38,41^. In contrast with ASJ, there are not promoters, to our knowledge, that drive the expression of genes exclusively in IL2L and/or PVQR neurons ^42^. We therefore used pairs of promoters, whose expression overlapped only in our neurons of interest, to drive expression of the two RNA strands, such that expression of dsRNA was restricted to IL2 or PVQ (see the Material and Methods section).

In agreement with our failure to rescue *unc-1(vlt10)* in ASJ, knock down of *unc-1* in ASJ did not induce any effect, supporting the idea that ASJ is not the site of function. In contrast, disruption of the expression of *unc-1* in IL2 neurons significantly increased polyQ aggregation, to the same level of *unc-1* mutants (Fig. 2D). Surprisingly, silencing *unc-1* in PVQ neurons induced a mild reduction of polyQ aggregation, suggesting again that gap junctions modulate polyQ aggregation (Fig. 2D). To confirm IL2 data, we reintroduced the cDNA of *unc-1* in IL2 neurons in *40Q*; *unc-1(vlt10)* animals, in two separate arrays each directing its expression from the *oig-1p* or the *osm-3p* promoters. These animals showed reduction of aggregation of polyQs in muscle cells and neurons (Fig. 2E, F). Altogether, these data confirmed that loss of function *unc-1(vlt10)* modulates polyQ aggregation, from IL2 neurons, which are electrically coupled to the ASJ neuron.

Our data are consistent with a model in which loss of *unc-1* function perturbs gap junctions between IL2 and ASJ, resulting in increased SSU-1-mediated signalling from ASJ. We wondered, therefore, whether the *unc-1* mutation resulted in an increase in general excitability of the ASJs. The ASJ neurons are sensitive to temperature. Previous studies showed that calcium concentrations in these neurons increased in response to warming and decreased in response to cooling ^43,44^. We therefore examined whether these responses to temperature were intact in *unc-1(vlt-10)* animals. We did not see any difference in the fluorescence ratio either at the starting temperature of 23 °C or after shifting to 17 °C (Fig. 2G-I), so tonic, global calcium concentrations do not appear to be increased. Indeed, when the temperature is shifted back to 23 °C we observe a defect in the up-step in calcium concentration (Fig. 2G, J-K), more suggestive, if anything, of a decrease in excitability. It seems more likely that these reflect distinct, localised functions.

### Ablation of the sulfotransferase gene *ssu-1* rescues *unc-1*-associated protein homeostasis unbalance

Carroll and co-workers showed that mutant alleles of *ssu-1*, a gene that encodes the only known alcohol cytosolic sulfotransferase in the genome of the worm, partially rescues uncoordinated locomotion of *unc-1* defective worms, among other phenotypes ^9^. Alcohol sulfotransferases are promiscuous enzymes that are able to add a sulphate moiety to a variety of molecules such as peptides, steroid hormones and xenobiotics targeted for elimination (reviewed by Hattori *et al*., ^45^). *ssu-1* expression is restricted to the ASJ pair of sensory neurons, which are involved in food sensing, hormone release, Dauer formation and temperature sensing ^9,43,46,47^.

To investigate the potential role of *ssu-1* as an interactor of *unc-1*, in regard to protein homeostasis, we introduced the insertion expressing *40Q* into worms carrying the loss-of-function alleles *ssu-1(fc73)* and *unc-1(e580). ssu-1* single mutant animals did not exhibit a modified aggregation phenotype (Fig. 3A). However, we found that mutations in *ssu-1* suppressed the effects caused by mutations in *unc-1* on protein aggregation in muscle cells (Fig. 3A). To confirm these results, we induced cell-specific RNAi against *ssu-1* in ASJ neurons of *unc-1(vlt10)* mutants, using the *trx-1* promoter and following a strategy described elsewhere ^38,41^. As expected, reducing the expression of *ssu-1* in ASJ partially rescued the aggregation phenotype of *unc-1(vlt10)* (Fig. 3B). These data indicate that SSU-1 activity in the ASJ sensory neurons is required for the disrupted protein homeostasis in muscle cells exhibited by *unc-1* mutants.

**Figure 3.**
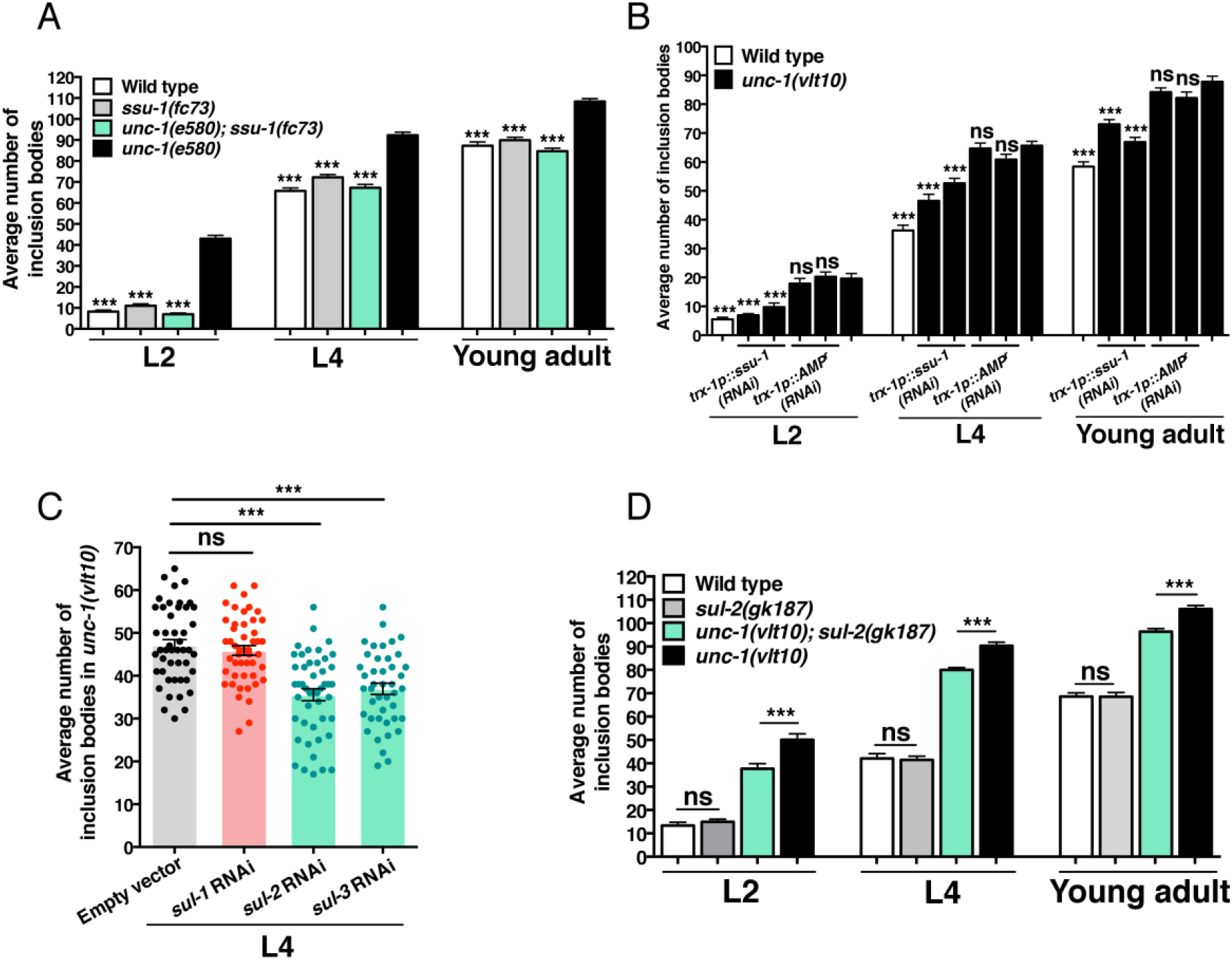
Neurohormonal signalling disruption produces an excess of signal that enhances polyQ aggregation. (**A**) Suppression of *ssu-1* does not modify the aggregation pattern of *40Q* animals, but rescues polyQ aggregation phenotype in *unc-1* mutants. (**B**) RNAi against *ssu-1* in ASJ neurons significantly reduces polyQ aggregation in *unc-1(vlt10)* worms. (**C**) Data represent the amount of polyQ aggregation after ubiquitous silencing of the different sulfatases of *C. elegans* (*sul-1, sul-2* and *sul-3*) into the *unc-1(vlt10)* background. RNAi against *sul-2* and *sul-3*, but not *sul-1*, reduces the polyQ aggregation pattern of *40Q*; *unc-1(vlt10)* compared controls. (**D**) Disruption of *sul-2* by the loss-of-function allele, *gk187*, in *unc-1(vlt10)* animals, partially reduces the average number of inclusion bodies in different stages (L2, L4, and young adult) while the simple mutant *40Q*; *sul-2(gk187)* shows no changes in aggregation phenotype at any stage. The plotted data show the mean ± SEM. For graphs A, B and D, 30 animals were analysed per condition and/or strain and per experiment. For graph C, 40 worms were analysed per condition and/or strain and per experiment. The analysis has been reproduced at least three times. ***: p-value < 0.001; ns: statistically not significant. The significance values are referred to *40Q*; *unc-1(vlt10)* strain per stage (graph A and B). Statistical test ANOVA with multiple comparative test type Tukey.

### Arylsulfatase activity modulates the aggregation dynamics of polyQs

Our results suggest that SSU-1 produces a sulfonated signalling molecule in *unc-1* mutants and that this sulfonated molecule promotes the aggregation of polyQs in muscle cells. Hence, we hypothesised that removal of sulfate in the signal may alter the aggregation phenotype somehow. To test this hypothesis, we sought to disrupt sulfatase enzymatic activity. There are only three genes encoding potential enzymes with predicted sulfatase activity in *C. elegans*: *sul-1, sul-2* and *sul-3* ^10^. *sul-1* encodes a predicted protein with similarity to mammalian 6-O-endosulfatases (www.wormbase.org; and Supplementary Fig. 4) and *sul-2* and *sul-3* predicted enzymes are closely related to arylsulfatases (www.wormbase.org; and Supplementary Fig. 4). Analysis of animals fed with bacteria expressing dsRNA targeting each of the three genes showed that reduction of function of both *sul-2* and *sul-3* decreased the aggregation observed in *unc-1* animals (Fig. 3C). In contrast, *sul-1(RNAi)* animals did not show a reduction of inclusion bodies (Fig. 3C), which suggests that only arylsulfatase function is able to modulate aggregation of polyQs in *unc-1* mutants. Then we analysed the aggregation phenotype of the only arylsulfatase mutant available, *sul-2(gk187)*, and we observed that ablation of *sul-2* reduces partially the aggregation phenotype of *unc-1* animals (Fig. 3D). These results support our findings obtained by RNAi, which suggests that arylsulfatase activity is required to modulate protein homeostasis.

### Activation of the nuclear hormone receptor DAF-12 reduces polyQ aggregation

Sulfotransferases such as SSU-1, add a sulfate moiety to steroid hormones, among other substrates. Moreover, some authors demonstrated that SUL-2, at least, is able to remove sulfate moiety from steroid hormones ^48^. Hence, we sought to investigate whether the signalling pathway of the dafachronic acids, may be produced in ASJ by disruption of *unc-1*, to disrupt protein homeostasis. Dafachronic acids induce opposing effects on longevity, depending on the genetic and metabolic context of the worms ^49,50^. The target of these hormones is DAF-12 ^51^, a well-studied nuclear receptor involved in longevity, reproductive development and lipid metabolism ^12–14^. To investigate whether DAF-12 modulates the phenotype of *40Q*; *unc-1* worms, we generated a lesion in *daf-12, vlt19*, using CRISPR, which affects the ligand binding domain (Supplementary Fig. 9B). However, homozygous *40Q*; *unc-1*; *daf-12* mutants were synthetic lethal, so we could not analyse them. In consonance with this, *40Q*; *daf-12* animals showed a substantial increase in the number of polyQ aggregates (Fig. 4A).

**Figure 4.**
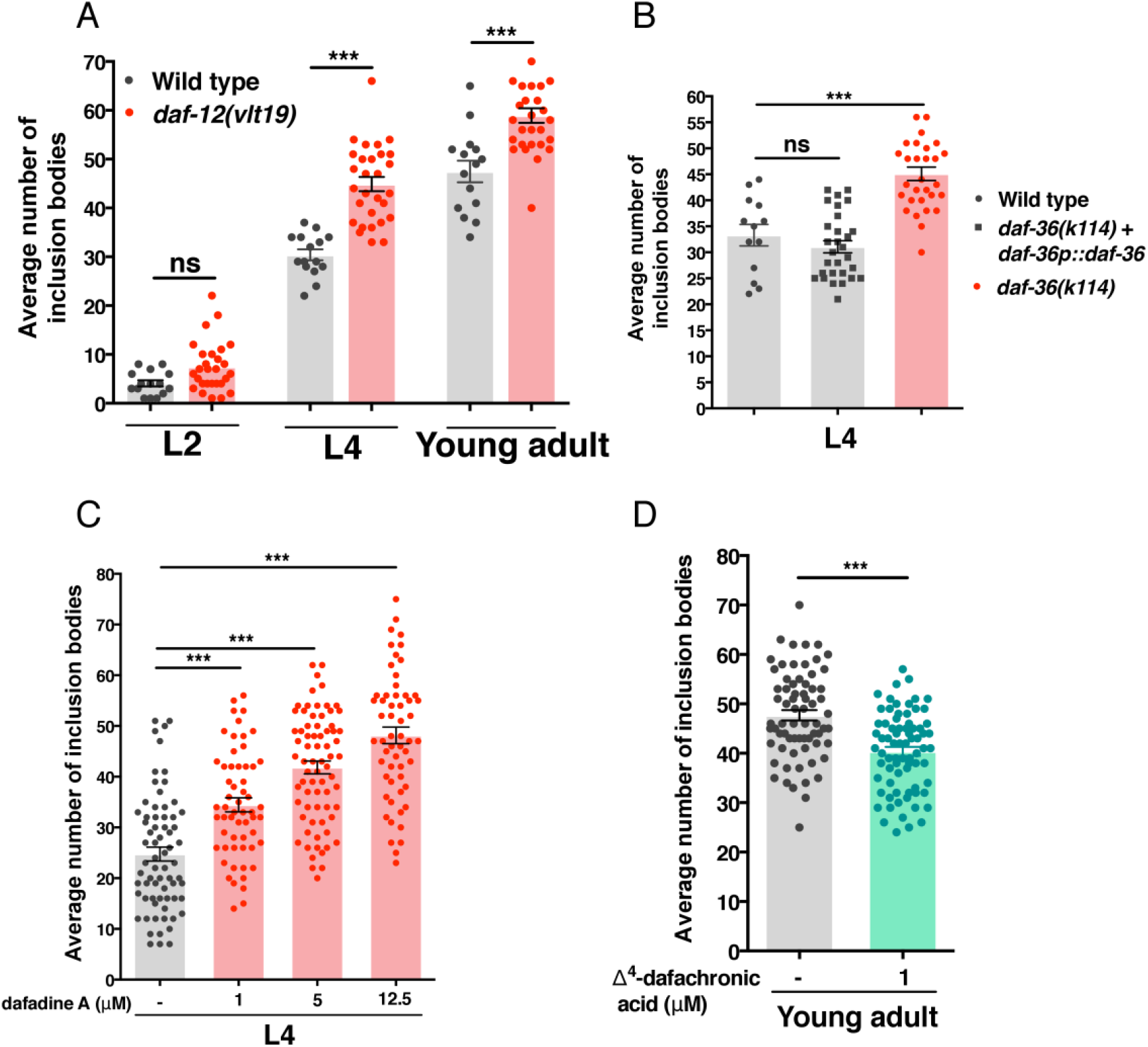
DAF-12 signalling is required to maintain protein homeostasis. (**A**) *daf-12(vlt19)* mutants show enhanced aggregation. (**B**) Suppression of *daf-36* increases formation of polyQ aggregation in wild type *40Q* worms and *daf-36(k114)* animals, compared with *daf-36* mutants rescued with an extrachromosomal array containing the whole genomic region of gene. (**C**) Blocking of *daf-9*, using dafadine A, shows a dose-dependent progressive increase of inclusion bodies in *40Q* worms. (**D**) Treatment of *40Q* with 1 μM Δ^4^-dafachronic acid reduces significantly the number of polyQ inclusion bodies in young adults. We analysed the following number of worms per strain and/or condition: for graph A and B 30, for graph C more than 55, and for graph D more than 65 worms. The analysis has been reproduced at least three times. ***: p-value < 0.001; ns: statistically not significant. The significance values are referred to untreated *40Q* animals (graph C, D). Statistical test ANOVA with multiple comparative test type Tukey (graph A, B and C). Statistical t-test with non-parametric Mann-Whitney test (graph D).

To further confirm these data, we investigated the effect of disrupting dafachronic acid biosynthesis, through inhibition of DAF-36 and DAF-9, two enzymes involved in their synthesis ^52^. The loss of function of *daf-36* allele *k114*, together with the insertion expressing *40Q*, was nearly sterile (data not shown). To avoid this issue, we rescued *daf-36* by reintroducing a DNA construct that contained the coding and regulatory region of *daf-36* into *40Q*; *daf-36(k114)* worms, as an extrachromosomal array. Then, we analysed the *daf-36* homozygous worms that lost the array. These animals showed a substantially higher number of inclusion bodies than their siblings carrying the array (Fig. 4B). To investigate the role of DAF-9 in the aggregation of polyQs, we used different concentrations of dafadine A (1, 5, 12.5 µM), a drug that specifically inhibits this enzyme ^53^. L4 *40Q* animals treated with this substance, showed similar polyQ-related phenotypes as *daf-12* mutants (Fig. 4C). Then, we analysed the behaviour of the *40Q* worms cultured on 1 µM Δ^4^-dafachronic acid, and we observed a reduced amount of polyQ aggregates (Fig. 4D). Collectively, these data suggest that signalling through DAF-12 is required to maintain protein homeostasis.

### Activation of the nuclear receptor NHR-1 induces enhanced polyQ aggregation

It is unlikely that DAF-12 is the target of the steroid signal from ASJ, because disruption of the pathway enhances aggregation, and potentially synergises with the pathway disrupted by *unc-1* loss of function. Hence, we sought to look at NHR-1, a nuclear receptor that regulates insulin sensitivity in *C. elegans* and which is controlled by SSU-1 ^54^. To introduce mutant alleles of *nhr-1* on *40Q; unc-1(vlt10)* worms we used CRISPR generating a lesion, *vlt16*, which emulates *nhr-1(n6242)* ^54^. Additionally, we isolated and characterized a new allele of *nhr-1, vlt15*, which was produced by an abnormal homologous recombination and which is a putative null allele, because it causes a change in frame in the gene and a truncated protein (Supplementary Fig. 9C). Although, we cannot rule out that this may be a neomorph allele, it is unlikely, because the allele needs to be in homozygosis to produce a phenotype (data not shown). Analysis of both mutant alleles showed that disruption of *nhr-1*, in *unc-1* worms, produces a substantial reduction of the inclusion bodies observed in *unc-1* mutants (Fig. 5A), indicating that NHR-1 acts downstream of SSU-1 to regulate protein homeostasis. In addition, *nhr-1* ablation also partially restored the uncoordinated phenotype of *unc-1* worms (Supplementary Fig. 5A). Reintroduction of *nhr-1*, by means of extrachromosomal arrays expressing its cDNA in body wall muscles, restored the aggregation phenotype shown by *unc-1* mutants while neuronal rescue did not (Fig. 5B). In addition, we show that *nhr-1* reintroduction aggravated motility to similar levels of *unc-1* mutants (Supplementary Fig. 5A). Rescue of the polyQ-induced motor defect is associated with a reduction of polyQ aggregation (Fig. 5B and Supplementary Fig. 5A). To further confirm this correlation between motor restoration and polyQ aggregation reduction, we treated *40Q* and *40Q; unc-1(vlt10)* animals with metformin, an anti-diabetic drug that reduces polyQ aggregation ^55,56^. As we expected, metformin treatment reduced polyQ aggregation and improves motor movement in *40Q* and *40Q; unc-1(vlt10)* animals, suggesting that motility rescue is due to decreased polyQ aggregation (Supplementary Fig. 5B, C).

**Figure 5.**
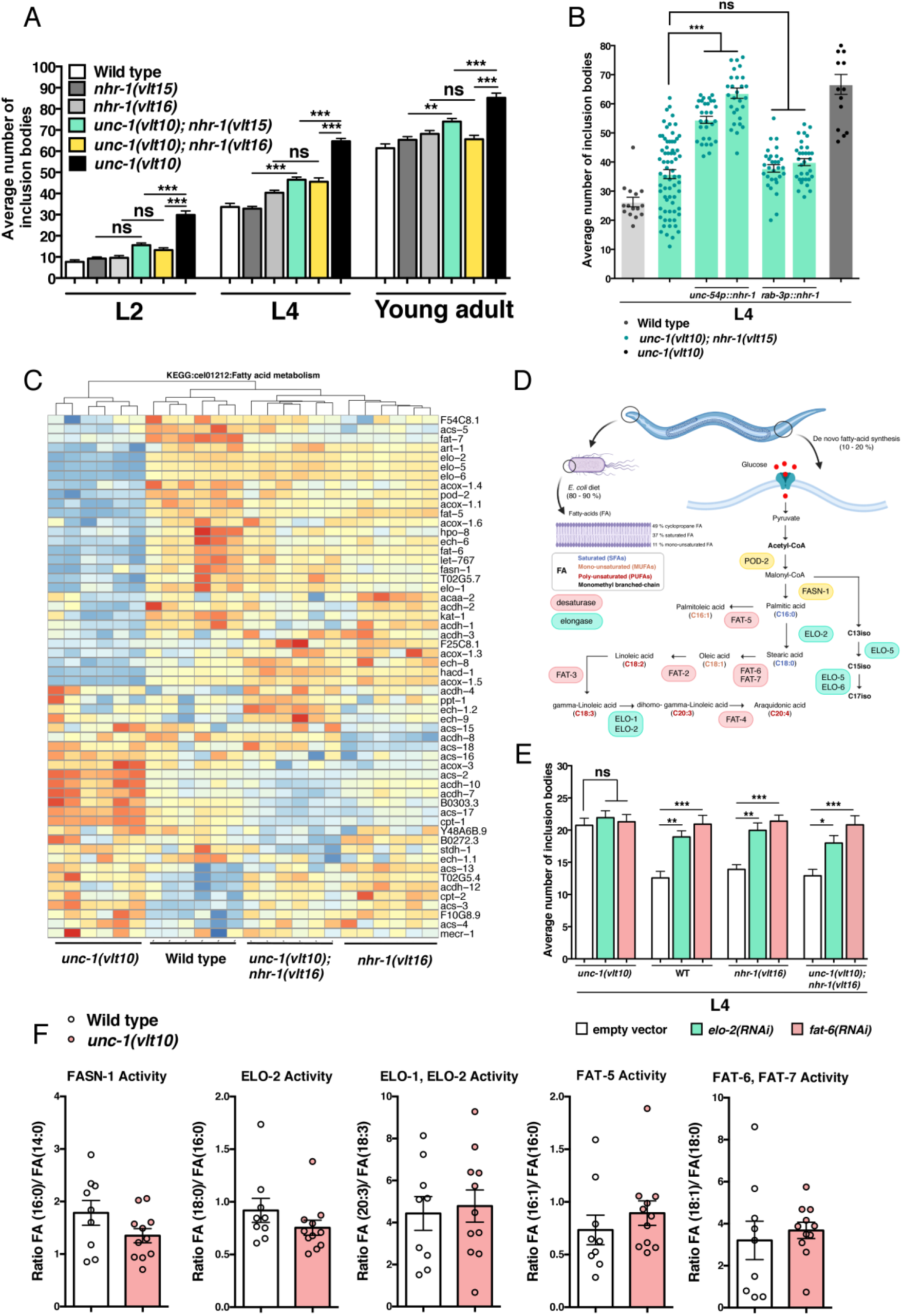
NHR-1 modulates protein homeostasis by controlling genes of the fat metabolism. (**A**) *unc-1* animals bearing *nhr-1* loss of function alleles *vlt15* and *vlt16* show dramatically reduction of aggregation of polyQs, while do no change proteostasis in *40Q* animals. *vlt15* phenocopies the *vlt16* allele, suggesting that both alleles are loss of function variants. (**B**) Reintroducing the cDNA of *nhr-1*, in muscle cells, increases the polyQ aggregation in the double mutant *unc-1; nhr-1*. In contrast, reintroduction of *nhr-1* in neurons do not modify aggregation of polyQs. (**C**) Heatmap showing the gene expression profile, related to the metabolism of fatty acids (KEGG: “Fatty Acid Metabolism”) of wild type, *unc-1, nhr-1* and double mutant worms expressing polyQs. Genes highlighted in blue show low expression levels while genes with high expression are shown in red in each biological sample included in the analysis. The genes that appear in the Heatmap have been selected according to a p-value < 0.05 and a fold change ≥ 2. The heatmap shows several genes of the synthesis of lipids, which are downregulated: *pod-2, elo-2, elo-5, fat-5, fat-6, fat-7*, among others, in *unc-1* mutants compared with the other genotypes. (**D**) De novo Fatty acid synthesis pathway diagram. (**E**) RNAi against *elo-2* and *fat-6* genes in different mutant backgrounds of *unc-1* and *nhr-1*. Ubiquitous silencing of *elo-2* and *fat-6* in *unc-1* mutants do not show a change of polyQ aggregation, while the number of inclusion bodies increases in wild type and *nhr-1* mutants. (**F**) Lipidomic assay in wild type and unc-1 mutants. Graphs show activities of some lipid enzymes FASN-1, ELO-1, ELO-2, FAT-5, FAT-6 and FAT-7. Ratio conversion of fatty acids (FA) is shown for each on lipid enzyme. We have performed lipidomic assay from around ten samples for each condition.

One of the features of polyQ aggregation is induction of some pathways of the unfolded protein response and genes encoding chaperones ^57,58^. Hence, we sought to test whether these pathways were switched on, in *unc-1* animals, and whether disruption of *nhr-1* may restore them. To do so we looked at expression of key genes involved in such signalling events. As expected, we observed that *unc-1* mutants show stress-ER and upregulation of UPR genes in ER, mitochondria and cytosol (Supplementary Fig. 5D; black bars). In contrast, ablation of *nhr-1* reduced the expression of some of these genes (Supplementary Fig. 5D; grey bars). Next, we sought to do an *in vivo* experiment, in which we tested the UPR^ER^ reporter, *hsp-4::GFP* ^59^, in wild type and *unc-1* mutants. The *vlt10* allele enhanced expression of this reporter, which further supports that *unc-1*; 40Q animals show higher ER-stress (Supplemental Fig. S5E, F). Altogether, these results show that *nhr-1* ablation restores a basal stress condition in *unc-1* mutants.

### NHR-1 regulates fat metabolism genes to modulate protein homeostasis

To elucidate the mechanism behind the effect of NHR-1 function, we performed transcriptomic analysis on the following strains: *40Q, 40Q*; *unc-1(vlt10), 40Q*; *nhr-1(vlt16)* and *40Q*; *unc-1(vlt10)*; *nhr-1(vlt16)*. This analysis showed that *unc-1(vlt10)* mutants exhibited a distinct transcriptomic signature compared to wild type, and that introducing the *nhr-1* mutation in *unc-1* mutants restored transcriptomic signature to wild type background (Supplemental Fig. S6). Among the altered genes, we identified differences in expression in genes encoding key enzymes of lipid metabolism (Fig. 5C, D). For example, we noticed that acetyl-CoA carboxylase (ACC) encoded by *pod-2* and fatty acid synthase *FAS* encoded by *fasn-1* show lower expression levels in *unc-1* mutants compared with double mutants (Fig. 5C). Both enzymes are involved in the first steps of *de novo* fatty acid synthesis. Elongases (*fat-5, fat-6* and *fat-7*), which catalyses carbon chain extensions of fatty acids also show altered expression in *unc-1* mutants, compared to the wild type worms (Fig. 5C). Other kind of enzymes, like desaturases, which are involved in removing hydrogen atoms from carbon and producing double bonds between them, are downregulated in *unc-1* mutants (Fig. 5C). In clear contrast, *nhr-1* ablation reverses the expression patterns of these genes, in *unc-1* mutants, and their expression is closer of that of wild type worms (Fig. 5C).

To validate the transcriptomic results, we analysed the role of the elongase *elo-2* and the desaturase *fat-6*, in regard of polyQ aggregation. Both genes are essential to produce oleic acid, a lipid which is neuroprotective in rodents ^60^, induce lifespan extension ^61^ and prevents polyQ aggregation ^21,22,62^ in worms. To validate the transcriptomic results, we used RNAi by feeding, to reduce their function, in wild type and mutant backgrounds (Fig. 5E). Silencing *fat-6* and *elo-2* expression increased the number of inclusion bodies in all strains, except in *40Q::YFP*; *unc-1(vlt10)* animals (Fig. 5E), in which there are already high levels of inclusion bodies. These functional data further suggest that NHR-1 controls the expression of some enzymes of the fat metabolism, which in turn are essential to maintain protein homeostasis.

### *unc-1* animals show altered fat metabolism and ablation of *nhr-1* restores their lipid profile

The production of oleic acid starts with ELO-2 converting palmitic acid (fatty acid chain with 16 carbons and no double bonds: C16:0) into stearic acid (C18:0), both of which are precursors of oleic acid. Then, FAT-6 cooperates with FAT-7 to synthesise oleic acid (C18:1) from stearic acid (Fig. 5D). Regarding this, we decided to investigate these enzyme activities to validate the transcriptomic data. We performed an untargeted lipidomic assay to find out the fatty acid content in wild type and *unc-1* mutants. This analysis showed that wild type animals shows a substantially different lipid profile (Supplementary Fig. 7). For example, whereas phosphatidylcholines are the most abundant lipids, in both strains, because they play a fundamental function as part of the structural membrane (Supplementary Fig. 7). However, the abundance in *unc-1* mutants is larger. In contrast, triglycerides and its precursors, fatty acids, correspond to 6-7 % each one of total in wild type animals, while these lipids represent around of 1 % of total in *unc-1* mutants (Supplementary Fig. 7). These results suggest that *vlt10* can induce a remodulation of lipid abundant by decreasing fatty acids and triglycerides content.

To better understand these results, we performed an analysis to study the activity of the enzymes that process fatty acids (FASN-1, ELO-1, ELO-2, FAT-5, FAT-6 and FAT-7) whose transcripts are downregulated in *unc-1* mutants. Fatty acid conversion can be, in principle, correlated directly with enzyme activity by measuring the ratio of product to substrate. In this regard, we calculated FASN-1 activity dividing the fatty acids with 16:0 per the fatty acids with 14:0, in wild type and *unc-1* worms (Fig. 5F). We observed a tendency that shows lower FASN-1 (ratio between 16:0 and 14:0 fatty acids) and ELO-2 activity (ratio between 18:0 and 16:0 fatty acids) in *unc-1* mutants compared to wild type worms (Fig. 5F). However, we did not observed differences according to the ratio between 20:3 and 18:3 fatty acids (measure of ELO-1 + ELO-2 activities) which suggests that ELO-2 may act on saturated fatty acid (upstream of the pathway; see Fig. 5D) conversion steps more than poly-unsaturated fatty acid (steps downstream; see Fig. 5D) processing in *unc-1* mutants. These results support our previous results that showed that *elo-2* silencing in double mutants, *unc-1; nhr-1*, restores polyQ aggregation in *unc-1* mutants. However, lipidomic analysis does not indicate a clear role fro FAT-6 since we do not see any differences on desaturase activity in *unc-1* mutants (Fig. 5F), in contrast with the polyQ aggregation effect associated to *fat-6* silencing in *unc-1; nhr-1* double mutants (Fig. 5E). As we mentioned before, we cannot control intake lipid from *E. coli*, which constitute around 80-90 % of lipid source and we cannot exclude that this influences our results. Nevertheless, altogether these results confirm that polyQ aggregation increase in *unc-1* mutants is caused by lipid metabolism alteration that is restored by *nhr-1* ablation.

## DICUSSION

Progressive malfunction of protein homeostasis is a natural characteristic of ageing in all organisms. Many pathways and molecules have been shown to influence this process, to maintain protein appropriately folded and located in the right cellular environment. Deleterious mutations in genes encoding components of some of these modulatory pathways accelerate the decline of protein homeostasis, and contribute to the progression of age-related disorders. Among these cellular processes, chemical synapses has been shown to control protein homeostasis in postsynaptic cells ^4^. In this work, we screened for genes that alter protein aggregation in worms expressing polyQs in body wall muscles. Through this screening, we identified the allele *unc-1(vlt10)* that enhances aggregation of aggregation-prone proteins (Fig. 1). *unc-1*, a *C. elegans* Stomatin-like protein encoding gene, regulates protein homeostasis in muscle cells and neurons, acting in IL2 neurons (Fig. 2), and aggregation of polyQs is specific of some stomatin-like proteins (Table 1). Loss of function alleles of several innexin genes produce a similar enhancement of aggregation, suggesting the involvement of gap junctions, which are the pore components of the electrical synapse, and other gap junctions, in invertebrates (see Dahl and Muller for a review,^63^. This suggests that electrical synapses are essential to maintain appropriate proteostasis in *C. elegans*. We found that loss of function of *unc-1* did not result in a general increase in ASJ excitability, as we would expect if ASJ-IL2 gap junctions simply served to shunt current. Rather, the differential effect on different ASJ functions suggests that these innexins play more complex roles, specific to different aspects of ASJ function. Innexins can also operate as hemichannels ^64,65^, so this is a potential explanation for this specificity.

Our results suggest that disruption of the communication between IL2 and ASJ, alters the function of the later, triggering an excess of secretion of sulfated hormones which is dependent on SSU-1 (Fig. 3). This excess of hormonal signal requires the function of SUL-2 and SUL-3 to induce an enhanced polyQ aggregation on muscle cells and neurons (Fig. 3), which suggest that the sulphur groups of this unknown hormone need to be removed. *sul-2* and *sul-3* are arylsulfatases, evolutionarily closer to human steroid sulfatases.

Our epistatic analysis shows that NHR-1 is the effector of the steroid hormone produced by ASJ, and that this nuclear receptor regulates lipid metabolism genes, which in turn have an effect in proteostasis in *C. elegans* (Fig. 5). Transcriptomic analysis of single and double mutants *unc-1* and *nhr-1* mutants shows that many genes of the fatty acid synthesis genes are altered in *unc-1* worms, and that introducing *nhr-1* loss of function alleles restores their expression close to wild type worms (Fig. 5). We have validated the role of the fat metabolism by reducing the function of *elo-2* and *fat-6* which increases polyQ aggregation (Fig. 5). FAT-6, together with FAT-7, convert stearic acid in oleic acid, a protective lipid, to promote longevity and improve neuronal proteostasis ^21,22^. We have shown that enzymes involved in oleic acid synthesis are downregulated in *unc-1* mutants; however, lipidomic data does not show changes in FAT-6 and FAT-7 activities. Regarding to this, we have to consider that we have calculated activity enzyme from ratios of lipids obtained from both de novo synthesis and diet, which could be interfering with expected results. Together all results suggest that fat metabolism alteration modifies polyQ aggregation, which involves a neurohormonal signalling (Fig. 6).

In contrast with NHR-1, the nuclear receptor DAF-12 is protective (Fig. 4). Disruption of this signalling pathway (i.e. mutation of *daf-12* or culturing worms on dafadine A, a well-known *daf-9* inhibitor) enhances polyQ aggregation, while addition of dafachronic acids ^66,67^ have a protective effect (Fig. 4). In agreement with our findings, Farina and co-workers showed that neurosteroids that activate DAF-12 are neuroprotective in *C. elegans* models polyQ-induced toxicity ^68^. Moreover, other authors showed that *daf-12* signalling promotes protein homeostasis, by modulating the function of the stress response of the ER ^69^. All these data strongly suggest that there are opposing effects, exerted by different steroid hormones and different signalling pathways, towards modulation of protein homeostasis. The wide range of metabolic processes regulated by nuclear receptors has made them outstanding druggable targets against inflammation, and metabolic and neurodegenerative diseases (see ^70–74^ for reviews).

Evidences indicate that neurosteroids may induce protection against neurodegenerative diseases that involves dysregulation of protein homeostasis, such as Alzheimer, Parkinson or Huntington diseases (see Borowicz *et al*., for a review (^75^). To our knowledge, this is the first report showing that different steroid hormone signals have opposing modulator effects in protein homeostasis in animals, and it will be of interest to investigate if this is evolutionary conserved between invertebrates and mammals. The function of some components of this pathway is amenable to be modified using drugs. For example, there are inhibitors against sulfotransferases ^76,77^, the cytochrome *daf-9*/ CYP2J2 ^78^, sulfatases ^79–81^ and modulators of NHRs ^82–86^, which are being investigated to treat different kinds of cancer and metabolic diseases. Some attempts has been made to use fatty acid-enriched diets to treat neurodegeneration, although the results were not very positive (see Bono-Yagüe et al., and Shanim et al., ^87,88^ for reviews). Probably, new approaches should focus on using modulators of key enzymes of the lipid metabolism, that alter the presence of protective lipids within the stressed tissue. Many of the pathways presented here, then, are potential targets to treat pathological conditions produced by toxic proteins, which are inducers of neurodegeneration. Finally, our work provides an *in vivo* genetically tractable model, *C. elegans*, to study these opposing hormonal signals and to screen for new therapies.

## MATERIAL AND METHODS

### Maintenance and culture of *C. elegans*

All strains were maintained under standard conditions ^89^. Some strains used in this work were obtained from Caenorhabditis Genetics Center (CGC; Minneapolis, MN, USA). The CGC is funded by the National Institutes of Health’s National Center for Research Resources (NGRR). CW911 strain: *ssu-1(fc73) V; unc-1(e580) X* was kindly provided by Phillip G. Morgan (University of Washington, Seattle, USA). Supplementary Table 1 provides a detailed list of strains used in this study. All strains were outcrossed at least three times and cultured at 20 °C.

### Genetic manipulation of *C. elegans*

To generate vectors for gene expression in *C. elegans*, we took advantage of Gateway^®^ MultiSite Pro system (Invitrogen, Waltham, MA, USA). Primer sequences for cloning are listed at Supplemental Table 2. Briefly, we amplified 1,7 kb from promoter region upstream of *rab-3* gene using specific primers wild type gDNA. We cloned the attB1-B2-flanked PCR product into pDONR^™^ 221-P1P5r to generate pDONR 221-P1P5r-*rab-3p* (pAPG1). We amplified the cDNA of *unc-1* to clone it into pDONR 221-P5P2-*unc-1* (pAPG2). We used pDONR 221-P1P5r-*myo-3p* kindly provided by Dra. Denisse Walker. Plasmids containing each promoter region, were recombined with the cDNA of *unc-1*, within a destination vector that contains the transcription terminator from the gene *unc-54*, pHP2 ^90^, to generate the following destination constructs: pDEST HP2-*rab-3p::unc-1::unc-54t* (pAPG3) and pDEST HP2-*myo-3p::unc-1::unc-54t* (pAPG4). To induce *nhr-1* expression in muscle cells and neurons we used pVD105 and pVD106 respectively ^54^.

To produce a dominant negative form of *unc-1*, the *n494* allele ^91^, we introduced the mutation into a cDNA of *unc-1* containing in pAPG2 to obtain pDONR 221 P5P2-*unc-1(n494)* (pAPG5). Next, we used pAPG1 and pAPG5 as donor vectors and pHP2 as destination vector, to generate pAPG6 (*rab-3p::unc-1(n494)::unc-54t*). To rescue *unc-1* in IL2 neurons we obtained fusion PCR products using *osm-3p* and *oig-1p* (see more detailed Supplementary Fig. 8). DNA mixtures for injection consisted of 25-50 ng/μL of the interest DNA [except for *myo-3p* or *unc-54p* expression analysis (1-5 ng/μL)], a DNA marker (pCFJ90; *myo-2p*::mCherry) at 2.5-5 ng/μL, together with the empty yeast plasmid pYES (Thermofisher, Waltham, MA, USA) as carrier DNA, at a total final concentration of 120 ng/μL. We injected each mix as described elsewhere ^92^.

### Cell-specific RNA interference

To induce tissue-specific RNAi we developed PCR fusion products, to produce constructs as described elsewhere ^38,41^. We fused a promoter region to sense and antisense fragments of the interest gene (Supplementary Fig. 8). Therefore, simultaneous expression of these DNA constructs in a specific tissue, such as nervous system (which are refractory to RNAi), induce a double complementary strand RNA synthesis to activate RNAi mechanism in neurons. We amplified promoter regions (of *rab-3, trx-1, oig-1, osm-3, glr-1, gpa-9*) to allow us to express dsRNA in a tissue-specific manner (nervous system, ASJ, IL2, PVQ neurons, respectively) (Supplementary Fig. 8 and Supplementary Table 2). In parallel, we amplified sense/antisense fragment from our genes of interest (*unc-1, unc-7, inx-2, inx-6* and *ssu-1*), adding a sequence complementary to the specific promoter that we wanted to fuse with it (Supplementary Fig. 8 and Supplementary Table 2). Both PCR products were combined in a single PCR reaction to amplify fused PCR products, using appropriate external primers complementary to the promoter and gene (Supplementary Fig. 8 and Supplementary Table 2). We used a DNA fragment from the bacterial β-Lactamase gene (which we named AMP^r^) as negative control in the tissue-specific RNAi experiments. We injected 2.5 ng/μL of each construct (sense/antisense) together with pCFJ90 (encodes a pharynx-expressed mCherry as a transgenic marker ^93^) as a reporter, and pYES (Thermofisher) as carrier DNA until 120 ng/μL. We injected each mix as described elsewhere ^92^.

### RNA interference by feeding

To induce ubiquitous and constitutive RNAi, we feed the animals with modified HT115 *E. coli* strain expressing dsRNA of each gene. We generated RNAi vectors for these genes cloning a coding fragment that covers all isoforms of each gene, into an EcoRV site of pL4440 vector. The PCR product for each gene was amplified using Phusion polymerase (Thermofisher) and specific primers (see detailed sequence in Supplemental Table 2). HT115 strains containing empty pL4440, as a control, and RNAi vectors (pAPG11-*sul-1*, pAPG12-*sul-2*, pAPG13-*sul-3*, pAPG52-*elo-2* and pAPG53-*fat-6*) were grown in liquid Luria-Bertani medium containing 50 µg/mL carbenicillin over night at 37°C. Next, bacteria culture was induced by IPTG for 2 h at 37°C, before seeding RNAi plates. After drying plates, synchronized animals (L1 or L3; depending of the assay) were feed with HT115 bacteria containing empty vector or RNAi vectors until reaching L4 stage to be scored.

### Construction of knock-in worms using CRISPR

We used CRISPR/Cas9 system to introduce point mutations into *nhr-1* and deletions into *inx-2* and *daf-12* sequences. We based on our design and strategy from Julián Ceron Lab (IDIBELL Institute, Barcelona, Spain) ^94^. To introduce deletions into *inx-2* and *daf-12* genes, we used two crRNAs for each one, which are complementary to distal exon regions. We generated a full deletion of *inx-2, vlt22*, using the following gRNAs: crRNA#1 5’ – ACCGGAGCTCTCCCACCAAA – 3’and crRNA#2 5’ – GCACAATGAGAAACCAGTAT – 3’. We used the ssODN sequence to isolate mutant strains easier: 5 ‘ – ACGTAGCGTTT GCGTGCGCACACCTCGCGTAGTGGTCCGCGTTCGCATTACTTGCGCTGGGGAAACATACTGTACTGATCGATCAAGAGTTTTCACTGTCTTCTCGTCCATCACCAGCCATATTCATAATTTCTTTCAAT – 3’ (neutral nucleotide are marked in yellow and it is useful to genotyping mutants) (Supplemental Fig. S9A). For *daf-12*, we generated *vlt19* allele using the following crRNAs: crRNA#1 5’ – ATATTATGGATGTTACCATG – 3’ and gRNA#2 5’ – GGAATCGTTGTTCGGAGAGC – 3’. *vlt19* encodes a deletion of 500 pb inside of ligand binding domain (Supplementary Fig. 9B). We targeted *nhr-1* to emulate *n6242* allele ^54^ by the following crRNA: 5’ – CACCACTCCACACCGTCTTC – 3’. The ssODN sequence to induce knock-in to generate the allele of interest was: 5’ – CCAACGAAGAAAATCAAGATGAGCAGCGGATCTGACGACGAGCAAGCCACCACTCCACACAGACTCTAAGACCAGGTCAGTGGGCGAAACACATTTACCCCAATTTGGATGCATCTTGAATTTCAACAA – 3’ to generate *vlt16* in *nhr-1* (in blue are marked neutral changes and in red are denoted the point-nonsense mutation (C/T) (Supplementary Fig. 9C). We isolated a new allele, *vlt15*, by an abnormal random recombination that encodes a premature stop codon later than *vlt16* (Supplementary Fig. 9C). To produce the mixture for injection, the crRNAs were suspended in 20 μL of IDTE nuclease free buffer to obtain 100 μM stock. ssODNs were suspended in the same buffer at 1 μg/μL. CRISPR components were added at the following final concentrations: Cas9: 4.5 μM; ALT-R trackRNA: 32 μM; target gene crRNA: 35 μM; ssODN target gene: 175 ng/μL. For targeting deletions, we used final concentrations 17.5 μM for each crRNAs. We used disruption of *dyp-10* gene as a selection marker as described ^95^.

### Quantitative PCR

The relative expression of genes was measured by RT-qPCR using the <u>ViiA7</u> thermal cycler from Applied Biosystems (Waltham, MA, USA) using Taqman™ probes. To verify the expression of genes included within extracromosomal arrays, we selected by hand worms expressing reporter (*myo-2p::mCherry*) for RNA extraction (OMEGA Bio-Tek, Norcross, GA, USA) using the M165FC dissecting microscope (Leica, Wetzlar, Germany). To evaluate expression of genes in mutants, we collected a synchronized population of young adult worms in RNA lysis buffer. Extracts were frozen at −80 °C before RNA extraction. RNA samples were treated by DNAase treatment (Qiagen, Hilden, Germany). cDNA synthesis was done using 0.1-1 μg approximately of RNA following the manufacturer protocol (Takara Bio, Kusatsu, Japan). We used the 2xPrimeTime^®^ Gene Expression Master Mix (Takara Bio), and the PCR program was as follows: 1 cycle of 10 min at 50 °C; 1 cycle of denaturalization at 95 °C 15 s; 40 cycles of polymerization at 60 °C 1 min. To normalize relative expression, we used the housekeeping control, *pmp-3* (Taqman probe from Applied biosystems, Ce02485188_m1). To measure expression of interest genes or transgenes we designed customized probes from IDTDNA. We considered all isoforms for each gene in the customized design. We included at least three technical replicates for each measuring.

### Analysis of the effect of Δ^4^-dafachronic acid and dafadine A on worms

Dafachronic acid treatment (Cayman chemical, Michigan, MI, USA) ^66,67^ was done as described ^15^. Briefly, we grown synchronized L1 animals on Dafachronic acid 1 µM (DA dissolved in ethanol) in M9 buffer containing 5 µg/mL cholesterol, 12.5 µg/mL nystatin, 50 µg/mL streptomycin and OP50 *E. coli* at 0.5 Optic density. Dafadine A (Sigma-Aldrich-Merck, St. Louis, MO, USA) was added at three dose (1, 5, 12.5 µM) using the same culture conditions ^53^. Animals were growth in liquid medium at 60 °C and moderate shaking until young adults. We evaluated the effect of both compounds at least three times in independent experiments over polyQ aggregation pattern.

### X-34 staining

X-34 dye (Sigma-Aldrich-Merck) stains specifically amyloid deposits as described elsewhere ^96^. One-day synchronized adults were incubated in a drop containing 1 mM X-34 and 10 mM TRIS pH 7.5 solution. Stained animals were washed with PBS-Tween and transferred to fresh NGM plates. Animals stained were mounting in agarose 2 X pads with a sodium azide drop (0,05 M) to count amyloid deposits by fluorescent microscopy (DM2500, Leica) and quantify stained area of worms by digital dissection microscope (DMD108, Leica).

### Quantification of polyQ inclusion bodies in muscle cells and neurons

*40Q::YFP* transgene induces body inclusion formation in muscle cells in an age-dependent manner ^25^. The average number of inclusion bodies were obtained by counting total number of polyQ::YFP inclusion bodies in muscle cells per animal *in vivo* using fluorescent microscope (M165FC, Leica). PolyQ aggregation produces a progressive phenotype with the age so we have reflected polyQ aggregation scoring in several stages to visualize this pattern (L2, L4 and young adult). Animals were analysed from heterogeneous population and after scoring, they were rejected to avoid undesired duplicates. Score assays were done, at least, in three independent experiments, in which we counted at least, ten animals per genotype and stage. Thirty animals or more were analysed for each genotype and stage in total.

To score neuronal polyQ aggregates we used a model that express 40 glutamines under the control of F25B3.3 promoter ^30^. Neuronal aggregation pattern is complicated to analyse in whole animals, so we selected Ventral Nerve Cord zone to show the average number of polyQ aggregates per genotype. Sixty young adults were analysed for each genotype in total after three independent scoring experiments. The number of neuronal polyQ aggregates was scored using DM2500 Leica vertical fluorescent microscope.

### Scoring of number of α-synuclein::YFP aggregates in muscle cells

The *unc-54p::α-synuclein::YFP* transgene induces late aggregation of the α-synuclein protein expressed in frame with the YFP fluorescent protein, specifically in muscle cells by *unc-54* promoter controlling ^32^. Unlike 40Q::YFP inclusion bodies, which are visible using a fluorescence microscope, α-synuclein::YFP aggregates are only detectable using high-resolution fluorescence microscopes (DM2500, Leica). Since it is not feasible to perform a complete count of whole muscle tissue of animals, we have selected the area located between the two pharyngeal bulbs to evaluate the aggregation pattern of α-synuclein, as described elsewhere ^97^ and using the DM2500 vertical fluorescence microscope (Leica). Sixty 2-day-old adults were analysed for each genotype in total after three independent scoring experiments. The number of α-synuclein aggregates in muscle cells was scored using DM2500 Leica vertical fluorescent microscope.

### Scoring of number of beta-amyloid X-34-stained deposits in muscle cells

To evaluate β-amyloid protein aggregation, we used a *C. elegans* model that expresses the human Aβ peptide constitutively in muscle cells ^33^. To detect β-amyloid deposits we stained 1-day-old adults with X-34 dye (Sigma-Aldrich-Merk). Stained 2-day-old animals were analysed by a vertical fluorescent microscope (DM2500, Leica). We selected the area between two pharyngeal bulbs to evaluate the number of β-amyloid deposits. Sixty 2-day-old adult animals were analysed for each genotype in total after three independent scoring experiments. The number of muscle β-amyloid aggregates was scored using DM2500 Leica vertical fluorescent microscope.

### Motility thrashing assay

We evaluated motility capacity of young adults by thrashing. This assay consists of scoring the number of thrashes that show an animal when swimming in M9 buffer. We have considered a thrash when animal moves simultaneous head and tail. Animals were acclimated for 30 sec before scoring number of thrashes for the following 30 sec. Each animal was collected independently in a well, avoiding its use after counting (pheromones could affect to animal behaviour). The average number of thrashes was extrapolated for 1 min to show the mean of the values. At least 15-20 animals were analysed for each genotype and transgenic line in three replicate times at 20 °C. Damaged animals during handling were discarded in this study.

### Transcriptomic analysis

L1 young adults 40Q::YFP, 40Q::YFP; *unc-1(vlt10)*, 40Q::YFP; *nhr-1(vlt16)* and 40Q::YFP; *unc-1(vlt10); nhr-1(vlt16)* animals were cultured in NGM plates at 20 °C until they reached the young adult stage. They were subsequently collected in M9 buffer, frozen, thawed and mechanically lysed with 200 mg of glass beads (Sigma-Aldrich-Merck) for 30 sec using a Fastprep shaker apparatus (Thermofisher Scientific, FP120 model). tRNA was extracted using NZY total RNA isolation kit, Nzytech. Supernatant was recovered from lysate samples by centrifuging extracts at maximum speed for 1 min. The purified RNA was treated with DNAse and quantified and sent for sequencing by Novogene (Cambridge, UK). We included six replicates for each strain.

Reads were aligned to *Caenorhabditis elegans* genome assembly WBCel235 using HISAT2 ^98^ was used for alignment, HTSeq ^99^ for gene expression quantification and DESeq2 ^100^ for differential expression analysis. Genes with a fold change greater than 2 and a corrected p-value lower than 0.05 were retained. R was used for downstream analysis ^101^, and the pheatmap package ^102^ for heatmap representations. Pathway analysis was performed using KEGG tool ^103^ (www.genome.jp/kegg/).

### Untargeted lipidomics assay

YA synchronized wild type and *unc-1(vlt10)* animals were collected in RIPA 1X buffer (Sigma-Aldrich-Merck) to obtain protein extract. We collected six biological samples per genotype in each independent experiment (n = 3 experiments and n = 10 cultures of worms per genotype). Total protein extract was quantified using the commercial kit (Pierce ™ BCA Protein Assay kit #23225, Thermofisher) to normalize the lipid abundance of the samples. Samples were analysed using a liquid chromatography equipment coupled to a high-resolution mass spectrometer with an orbitrap detector (UPLC-Q-Exactive Plus) and an electrospray source (ESI) following the procedures optimized in the Analytical Unit (internal method not published). Samples and quality controls were randomly injected into the chromatographic system to avoid variability within the analytical sequence, as well as to improve the quality and reproducibility of the study. Data were acquired in in Full MS, DIA (data independent analysis) and DDA (data dependent analysis) scan modes and processed using an in-house script in the R software (v.3.6.1) with the XCMS and CAMERA packages for detection, filtering and alignment of spikes. The LipidMS library was used to identify candidate lipids ^104^.

### Imaging worms by microscopy

Fluorescence images were acquired using an SP5 confocal microscope and DM2500 vertical fluorescent microscope (Leica). Oil Red O images to analyse stained area per animal were taken with a digital dissecting microscope DMD108 (Leica). In all cases, animals were mounted on 2X agarose agar surfaces containing one drop of sodium acid (0.05 M) to anesthetize them. Nematode selection and manipulation was performed using dissecting microscopes without or with fluorescence (MS5 and M165FC, Leica).

### Statistics

Statistical analysis was performed using an analysis of variance test (ANOVA) combined with a Tukey test to perform multiple comparisons of different strains and/or conditions. To perform comparisons between two conditions we used a Mann Whitney t-test to obtain the statistical significance of the data. In the graphs we show the mean ± standard error (SEM). SEM is displayed as a bar, while asterisks show significant of the data (p-value) (* <0.05; ** <0.01; *** <0.001). Abbreviation “ns” denotes not significant differences.

## ACKNOWLEDGMENTS

We wish to thank the CGC (funded by NIH Office of Research Infrastructure Programs; ref: P40 OD010440) and Phillip Morgan for worm strains. We thank the Microscopy Unit and the Genomics Unit of IIS-La Fe, for their kind help. We also would like to thank Julián Cerón, for sharing protocols to induce CRISPR. We are grateful to Sara Torres and Howard Baylis for help and plasmids. We also thank AVAEH, the Valencian Association of the Huntington Disease, for funding part of this work. RPVM held a “Miguel Servet” fellowship (Ref: CPII16/00004) funded by the Instituto de Salud Carlos III (ISCIII, Madrid, Spain) and by Social European Funds, and grants from the ISCIII (PI17/00011and PI20/00114). The funds from the ISCIII are partially supported by the European Regional Development Fund. RVM also received an Ayuda Miguel Gil grant to RPVM (VII Convocatoria Ayudas a la Investigación MHER, 2019, cofinanced by Colegio Oficial de Farmacéuticos de Sevilla and Fundación Cajasol).

## AUTHOR CONTRIBUTION

APG-E and RPV-M designed the study, performed most experiments, analysed the data and wrote the manuscript. CM, JMM, RS and JK contributed with essential reagents and ideas and wrote the manuscript. MR and AL performed and analysed metabolomics data and wrote the manuscript. JP and CM analysed the data from RNA sequencing and wrote the manuscript. QC-Z, IA-R and JB-Y performed some experiments. JB and JC analysed high-throughput to isolate the *unc-1(vlt10)* mutation. NB provided essential reagents and wrote the manuscript.

