## Supplementary material for "Opposing steroid signals modulate protein homeostasis through deep changes in fat metabolism in *Caenorhabditis elegans*"

### SUPPLEMENTARY RESULTS

#### Isolation of the *unc-1(vlt10)* mutation

We induced random mutagenesis using 47 mM EMS (methanesulfonic acid ethyl ester, Sigma, St. Louis, Missouri, USA) in L4 animals of the AM141 strain. We incubated worms for 4h at 20°C on this solution. After washing them they were pipetted onto NGM plates, seeded with OP50, and allowed them to lay the F1. F1 animals were bleached when they reached adulthood, and we searched among the F2 for animals uncoordinated and with abnormal aggregation patterns. Once isolated animals with the right phenotype, they were outcrossed 5 times against the wild type background (N2, Bristol), before high-throughput sequencing of their genomic DNA. The reads provided by the sequencing service (Centre Nacional d'Anàlisi Genòmica – Centre de Regulacio Genomica, Barcelona, Spain) were mapped against the *C. elegans* WS245 reference using the mem algorithm implemented by the BWA software [1,2]. BAQ qualities were calculated and applied to the BAM alignments by using samtools calmd [1]. The SNP calling process was carried out by Freebayes [3] with a minimum mapping quality of 57, a base quality threshold of 20, a minimum coverage of 6 and a minimum SNP quality of 20. A filtering process was stablished to look for the SNPs whose allelic frequencies were likely to have been affected by the selection process. SNPs were filtered out if they had more than one allele in all families, or if the reference allele was not present in any family, or if there were more than three mutant alleles when all families were considered. For the SNPs that passed all the filters a selection index was calculated. It consisted in the difference between the frequency of the most frequent allele in the back-crossed population and the frequency of that same allele in parental population, thus the SNPs with the highest differences between both populations would had the highest selection index. The predicted effect of each SNP was calculated by SnpEff [4].

#### Selective depletion of the electrical synapse modulates polyQ aggregation

Besides *unc-1*, there are other nine genes that encode stomatin-like protein in *C. elegans* [5]. As *unc-1* functions in neurons, we tested whether some stomatin-like proteins, expressed exclusively in neurons, modulate polyQ aggregation in muscle cells. The genome of *C. elegans* contains four neuronal genes encoding stomatin-like proteins, *mec-2*, *unc-24*, *sto-1* and *sto-6* [6,7] Analysis of two of them genes, *mec-2* (expressed in mechanosensory neurons) and *unc-24* (wide neuronal expression), showed that only *unc-24* mutants have a phenotype comparable to *unc-1* mutants, suggesting that not all neuronal stomatin-like proteins are involved in the regulation of polyQ aggregation (Table 1).

Since *unc-1* modulates electrical synapse and gap junctions [8], we tested whether innexins (gap junction components) are polyQ aggregation modulators. To test this hypothesis, we introduced mutant alleles of some innexins widely expressed in the nervous system (*unc-9* and *inx-7*) and other genes that are expressed in a restricted set of neurons (*inx-2*, *inx-6* and *unc-7*), into a polyQ background [9] (Table 1). However, we observed a selective effect where only the ablation of *unc-7*, *unc-9* and *inx-2* enhances polyQ aggregation similar to *unc-1* mutants (Table 1).

The data shown above show that disruption of some components of the gap connections alters polyQ aggregation. However, this does not show that the neuronal electrical synapse *per se* is involved in regulating protein homeostasis. To confirm our results, we performed RNAi against innexins *inx-2* and *unc-7* are expressed in a restricted set of neurons [9]. Silencing of *inx-2(RNAi)* and *unc-7(RNAi)* increased inclusion body formation, in contrast with *inx-6(RNAi)* that did not modify aggregation pattern (Supplementary Fig. 3A). Then, we investigated whether *unc-1* interacts with *inx-2* or *unc-7*. The double mutant *40Q*; *unc-1(vlt10); unc-7(e5)* produce aggregation to similar levels than simple *unc-1* mutants (Supplementary Fig. 3B), which suggest that both molecules may work in the same process. In contrast, *40Q*; *unc-1(vlt10); inx-2(vlt22)* animals showed a clear additive effect suggesting that *unc-1* and *inx-2* operate in parallel pathways to modulate protein homeostasis (Supplementary Fig. 3C). Altogether, these results suggest that neuronal electrical synapse modulate protein homeostasis.

**Reduction of polyQ aggregation in muscle cells translates in better motor behaviour**

Carroll and co-workers showed that disruption of *ssu-1* rescued the uncoordinated phenotype of *unc-1* mutants [6]. In this regard, disruption of NHR-1 induced a similar rescue of motility in *40Q*; *unc-1; nhr-1* mutants (Supplementary Fig. 5A). In contrast, reintroduction of NHR-1 function in muscle cells, in the *40Q*; *unc-1; nhr-1* worms, substantially reduced the motility of the worms (Supplementary Fig. 5A). These results suggest that *nhr-1* mutations phenocopy the *ssu-1* mutations in *unc-1* mutants. To confirm that the rescue of the movement is associated to the modulation of protein homeostasis, and not to synaptic activity regulation of *unc-1*, we decided to investigate whether the reduction of inclusion bodies was responsible of this effect. In this regard, it is well-known that 2 mM metformin activates AMPK to reduce polyQ inclusion bodies in *40Q* animals, and that this leads to an improvement of motor behaviour [10,11]. Hence, we treated *40Q*; *unc-1(vlt10)* worms with metformin, which reduced the average of body inclusion formation (Supplementary Fig. 5B), and also improved the movement defect in both *40Q* and *40Q*; *unc-1(vlt10)* animals (Supplementary Fig. 5C).

***unc-1* animals show stress in the ER which is rescued by ablation of *nhr-1***

We hypothesised that the enhanced polyQ aggregation in *unc-1* animals may activate unfolding protein response pathways (UPR). If that was the case, disruption of *nhr-1* should reduce the associated stress in *unc-1(vlt10)* worms, and deactivate these pathways. Hence, we performed real-time PCR in some genes related to the UPR of the mitochondria ($UPR^{mt}$), endoplasmic reticulum ($UPR^{ER}$) and cytosol (HSR), in *unc-1* and *nhr-1* mutants. As expected, *unc-1* mutants showed an increased expression of some of these genes, while double mutants *unc-1; nhr-1* showed reduced expression, which suggest that they have less stress (Supplementary Fig. 5D).

To further demonstrate this, we introduced the *vlt10* mutation into a ER-stress reporter, *hsp-4::GFP*. To increase the ER-stress signal we treated *unc-1(vlt10)* animals with mild amounts of tunicamycin (see the Material and Methods section) and compared them with wild type worms. After treatment, *unc-1(vlt10)* animals showed higher *hsp-4::GFP* expression levels than wild type animals which further shows that *vlt10* animals have ER-stress (Supplementary Fig. 5E, F).

**Transcriptome analysis shows several cellular processes altered in *unc-1* mutants that recover after ablation of *nhr-1***

Among the many genes which expression was altered in *unc-1(vlt10)* mutants, we have identified 532 genes that are rescued by the *nhr-1(vlt16)* mutation (Supplemental Fig. 6D). To identify cellular processes that where altered in *unc-1* mutants, we used the WormBase tool for Gene ontology (GO) analysis [12,13] and KEGG pathway analysis [14]. We found that genes that were differentially expressed in <u>*unc-1* mutants were significantly enriched in terms related to</u> lipid metabolism, including fatty acid metabolism (KEGG:cel01212), biosynthesis (KEGG:cel00061 ), degradation (KEGG:cel00071 ) and fat content increase (WBPhenotype:0001184), UPR response and immune response, among others. In addition, some lipid genes that show downregulated expression in *unc-1* worms, are restored in *unc-1; nhr-1* mutant animals (Fig. 5C). This suggests a direct link between regulation of lipid metabolism by *unc-1* and *nhr-1* and polyQ aggregation.

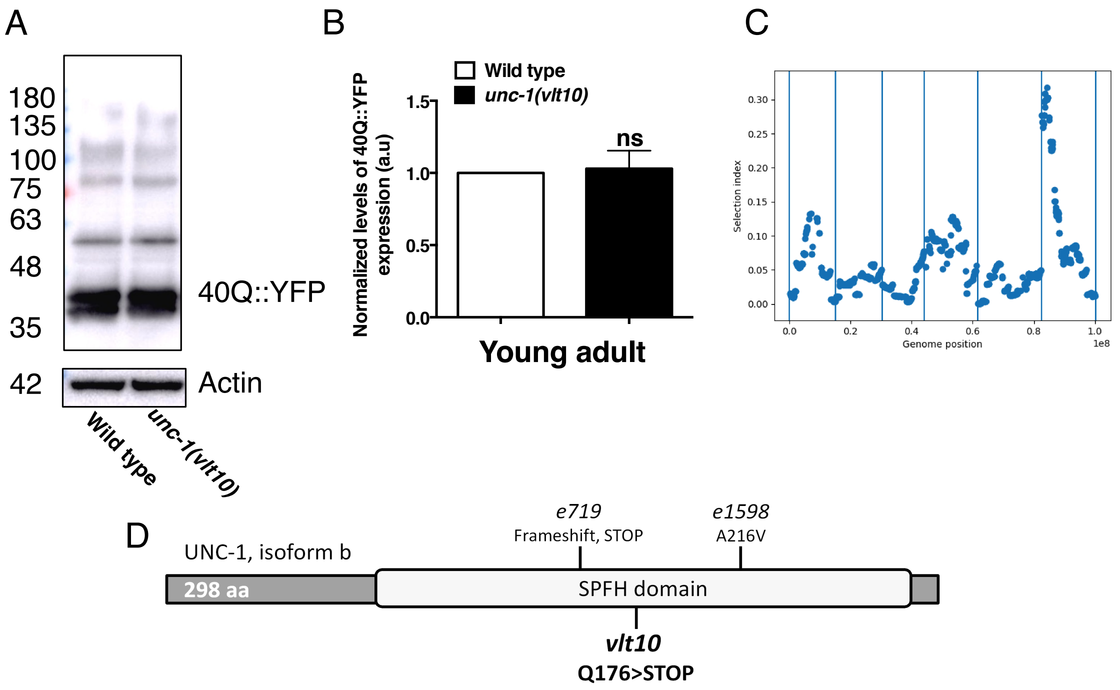


**Supplementary Figure 1. *vlt10* allele produces a premature stop codon in *unc-1/Stoml3* which enhances polyQ aggregation.** (**A-B**) Western blot shows that the expression of polyQs do not change in *unc-1* mutants. (**C**) A map of SNPs which suggest that *vlt10* is located into a small region of the X chromosome of the *C. elegans* genome. (**D**) *vlt10* produces a premature stop codon (Q176> STOP) in *unc-1*, in the SPFH domain, which likely produces a truncated UNC-1, and a putative loss of function.

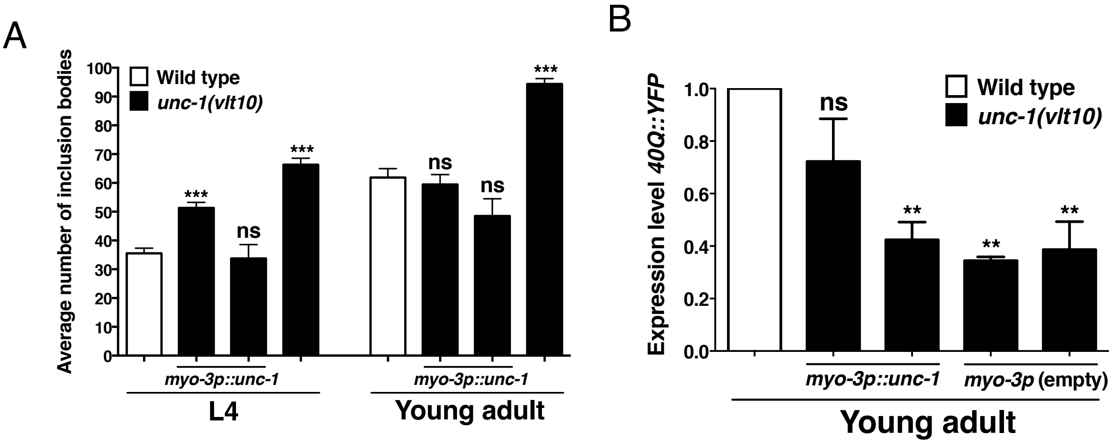


**Supplementary Figure 2**. **Reintroduction of *unc-1* in muscle cells of *vlt10* mutants reduces *40Q* expression, due to a change in polyQ expression.** (**A**) Introduction of a construct containing the cDNA of *unc-1* under the control of the promoter of the *myo-3* gene, in *unc-1* mutants, decreases the number of inclusion bodies differentially in L4 larvae and young adults. (**B**) Relative expression levels of the *unc-54p::40Q::YFP* transgene are substantially reduced in both transgenic lines expressing the *myo-3p::unc-1(cDNA)* transgene. Strains carrying arrays containing just the *myo-3p* promoter show a substantial reduction of the polyQ aggregates, suggesting that this promoter alters the expression of the polyQ transgene. YFP levels are normalized using *pmp-3* as a housekeeping gene, and relative to the control strain *40Q*. The plotted data show mean ± SEM. We used at least 30 animals per experiment and condition and we performed at least three independent experiments. ***: p-value < 0.001; **: p-value < 0.01; ns: statistically not significant. Statistical test ANOVA with Tukey's posthoc analysis.

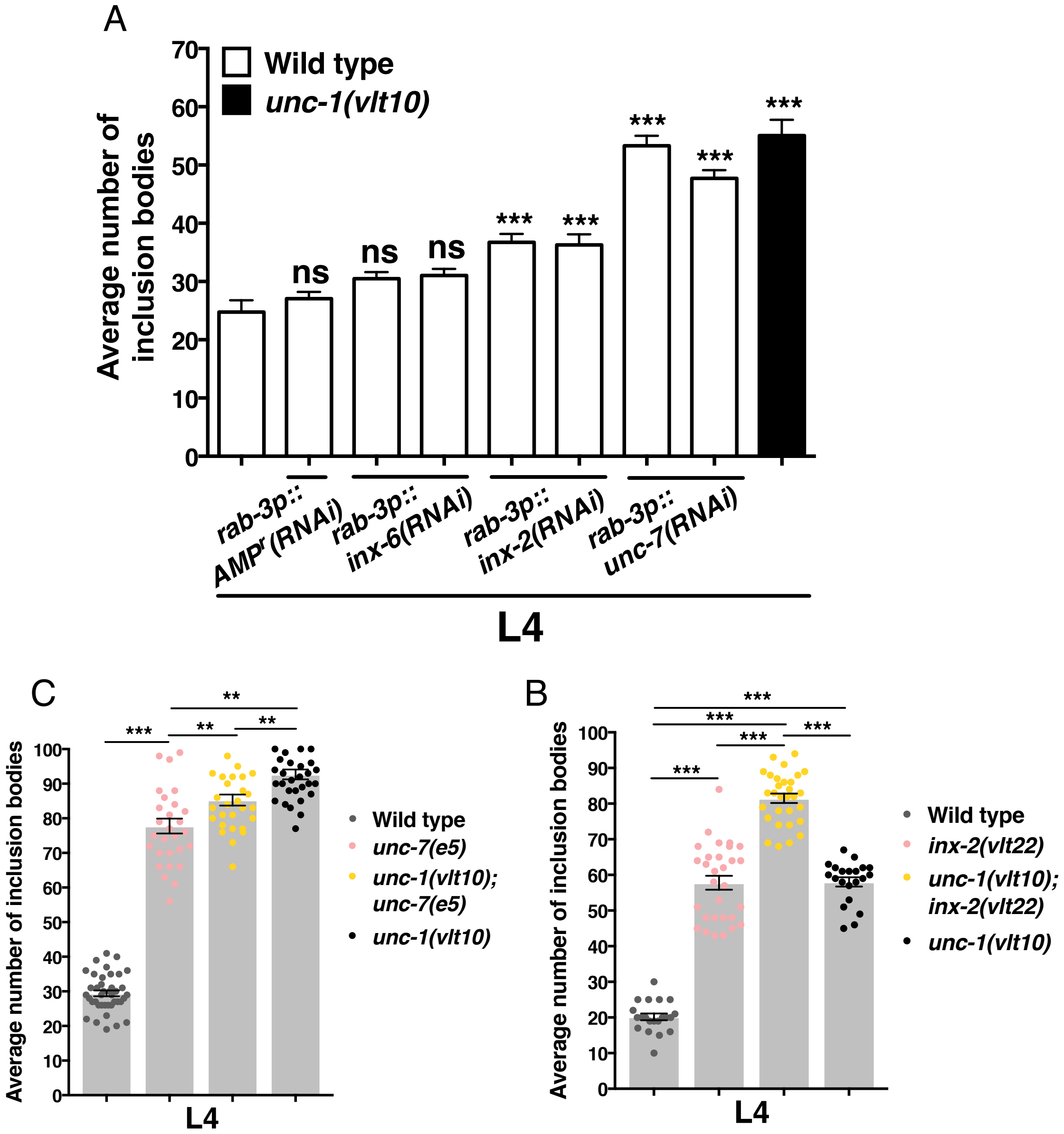


**Supplementary Figure 3. Neuronal disruption of innexins INX-2 and UNC-7 enhance aggregation of polyQs in muscle cells.** (**A**) Tissue-specific silencing of *inx-2*, *inx-6* and *unc-7* in neurons show a different effect over polyQ aggregation. RNAi against *inx-2* and *unc-7* induce increased polyQ aggregation in contrast to *inx-6* silencing. (**B**) Double mutant *unc-1(vlt10); inx-2(vlt22)* show the highest average number of polyQ inclusion bodies compared with simple mutants and wild type animals, which suggest that both mutations, *vlt10* and *vlt22*, have an additive effect on the aggregation phenotype. (**C**) *unc-7(e5)* and *unc-1(vlt10)* increase polyQ aggregation independently. The combination of both alleles into a polyQ context induce an intermediate average of the number of polyQ inclusion bodies compared to simple mutants, which suggests that *unc-7 (e5)* and *unc-1(vlt10)* could be working in the same process. The plotted data show mean ± SEM. 30 animals were analysed per condition and/or strain and per experiment. The analysis has been reproduced at least three times. ***: p-value < 0.001; **: p-value < 0.01; ns: statistically not significant. Significance p-values are referred to the wild type *40Q* strain (graph A). Statistical test ANOVA with multiple comparative test type Tukey.

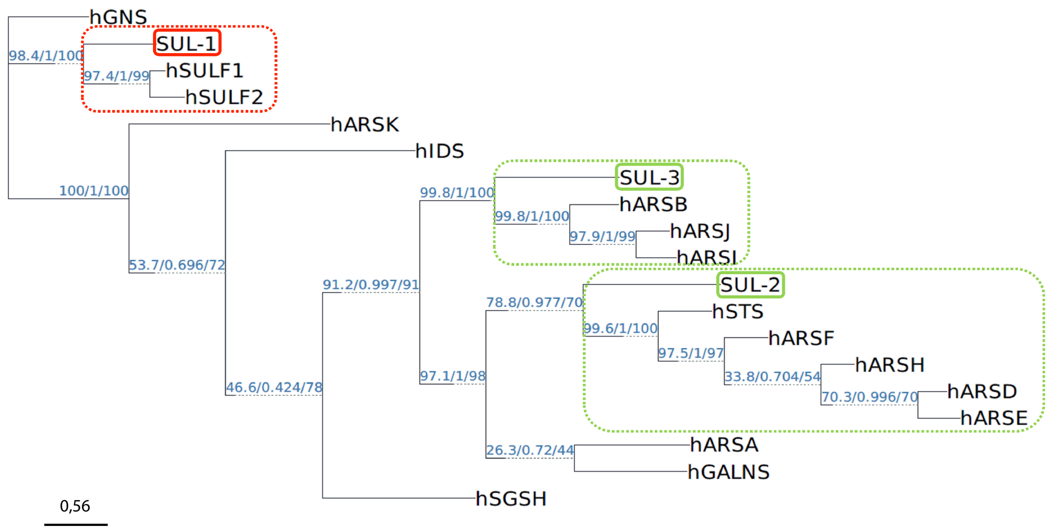


**Supplementary Figure 4. Phylogenetic analysis of *C. elegans* and human sulfatases.** The diagram shows the phylogenetic tree of all *C. elegans* sulfatases (SUL-1, SUL-2 and SUL-3) obtained using the MUSCLE (Edgar, 2004) and IQ-TREE version 1.6.8 software [2,15,16]. The phylogenetic tree shows bootstrap values which provides confidence values for each node. This analysis suggests that SUL-2 and SUL-3 are closer to arylsulfatases, while SUL-1 is closer to hSULF1 and hSULF2.

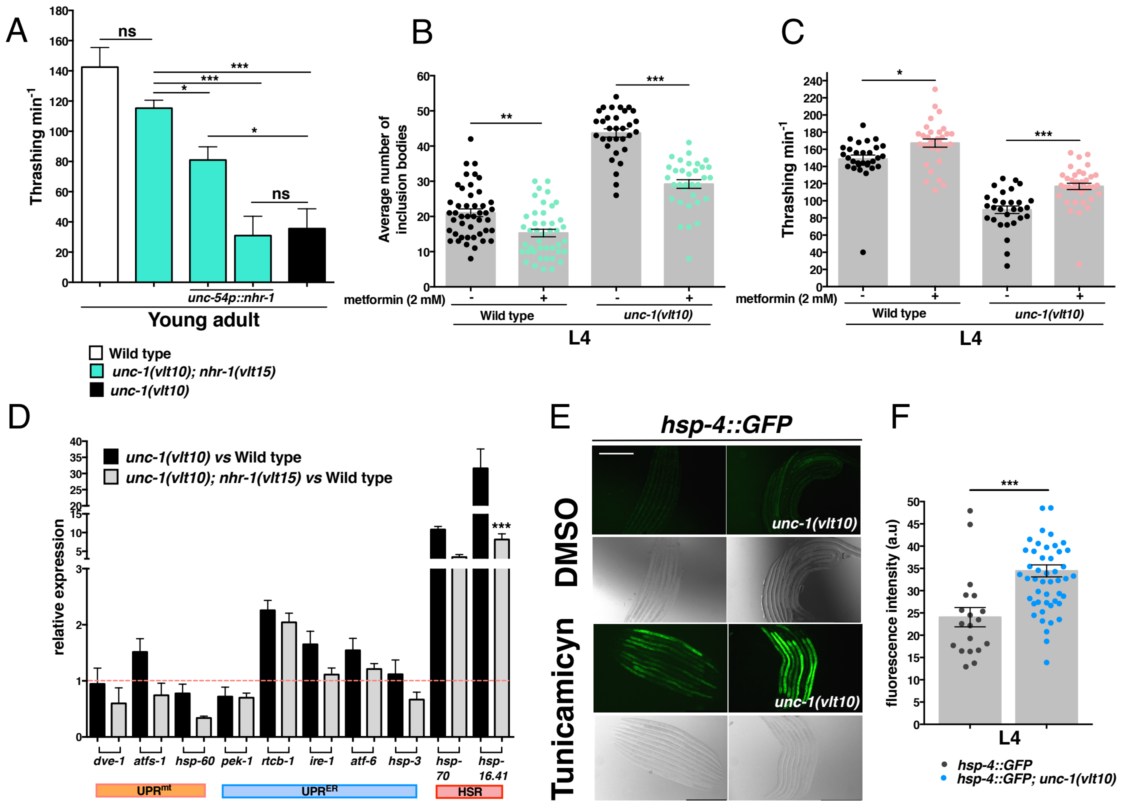


**Supplementary Figure 5. Ablating *nhr-1* improves movement and health span and reduces *unc-1(vlt10)*-associated stress.** (**A**) muscle-specific restoration of NHR-1 induce a lower motility capacity of *unc-1(vlt10); nhr-1(vlt16)* young adults compared to non-rescued double mutants. (**B - C**) Metformin treatment reduces polyQ aggregation and improves health span of wild type and *unc-1* mutants. (**D**) *unc-1* disruption induces stress that increases the expression of several Unfolded Protein Response (UPR) genes in the endoplasmic reticulum, cytosol and mitochondria compared to wild type (discontinuous red line). Some UPR-related genes are downregulated in *nhr-1* mutants compared to *unc-1* animals. (**E**) Representative images from wild type and *unc-1(vlt10)* animals that expresses the *hsp-4::GFP* transgene. The expression of this transgene is activated by a mild treatment with tunicamycin (1 µg/mL), which induces stress in the endoplasmic reticulum. (**F**) The loss of *unc-1* function induces higher expression of the reporter.

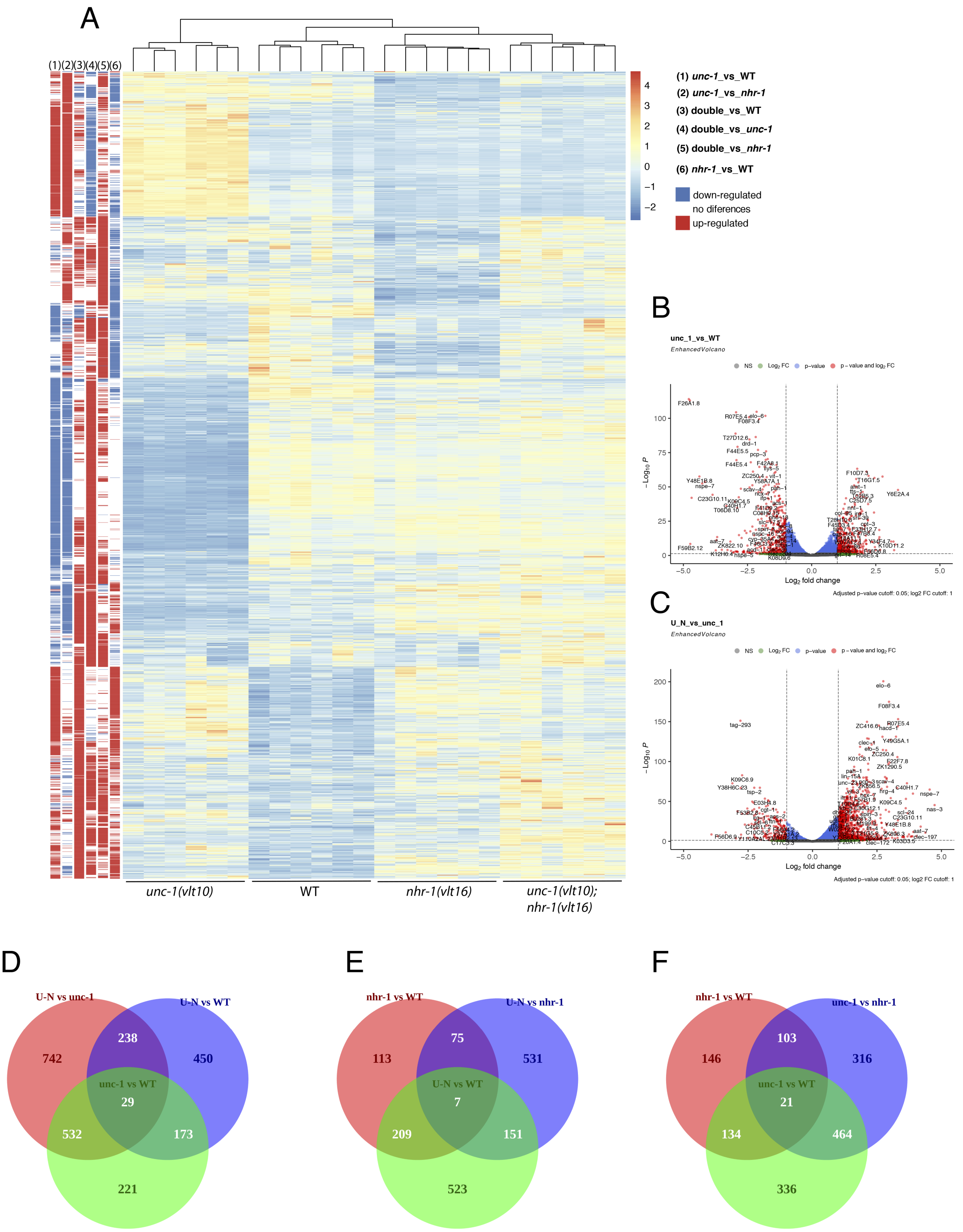


**Supplementary Figure 6. Transcriptomic signature of *unc-1* and *nhr-1* mutants. (A)** Heatmap showing DESeq2 normalized read counts of differentially expressed genes (corrected p-value < 0.05 and fold change > 2, see Methods). Data were scaled and centered by gene. Genes which expression level was rescued in *unc-1* mutants by the *vlt16* allele, have expression levels similar to both wild type (WT) animals and *nhr-1* simple mutants. On the left side of the heatmap, statistical significance of the different comparisons between genotypes have been included (denoted as (#1; *unc-1(vlt10)* vs WT), (#2; *unc-1(vlt10)* vs *nhr-1(vlt16)*), (#3; double mutant vs WT), (#4; double mutant vs *unc-1(vlt10)*), (#5; double mutant vs *nhr-1(vlt16)* and (#6; *nhr-1(vlt16)* vs WT). **(B-C)** Volcano plots showing Log2 Fold Change in the x axis versus -Log10 of the DESeq2 corrected p-value in the y axis, for each pair of genotypes. **(D-F)** Venn diagrams show the number of genes that are differentially expressed between genotypes. For example, graph D shows that there are 523 genes that are differentially expressed between double and *unc-1* mutants, and also between *unc-1* and WT, but not between double mutants and WT , reflecting that these genes are rescued by the *vlt16* allele. In contrast, there are 173 genes differentially expressed that are specific of animals carrying the *vlt10* allele and are not altered by *vlt16*.

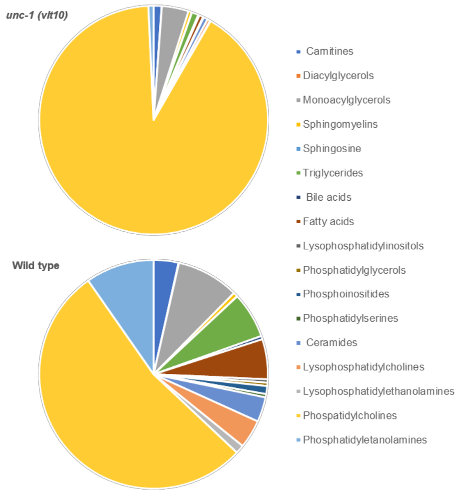


**Supplementary Figure 7**. Distribution of lipid classes in *unc-1(vlt10)* and wild type animals.

### SUPPLEMENTARY MATERIAL AND METHODS

#### RNA sequencing and analysis

Extracted RNA was sent to NOVOGENE (Cambridge, UK) for library preparation and sequencing. Six biological replicates, consisting of pooled bulk nematode RNA were sequenced for each genotype. Briefly, after quality control, mRNA was enriched using oligo(dT) beads and randomly fragmented. cDNA was synthesized using random hexamers and reverse transcriptase. After first-strand synthesis, a custom second-strand synthesis buffer (Illumina) was added with dNTPs, RNase H and *Escherichia coli* polymerase I in order to generate the second strand by nick-translation. To prepare the final cDNA library, a round of purification, terminal repair, A-tailing, ligation of sequencing adapters, size selection and PCR enrichment were performed. Library concentration was first quantified using a Qubit 2.0 fluorometer (Life Technologies). Insert size was checked on an Agilent 2100 and quantified using quantitative PCR (Q-PCR). After sequencing, reads with adapter contamination, with uncertain nucleotides in more than

10% positions or with a Phred score < 20 in more than 50% of the nucleotides were removed. Clean reads were aligned to *Caenorhabditis elegans* genome assembly WBCel235 using HISAT2 [17], and gene expression was quantified with HTSeq [18] using the union model. DESeq2 with default options was used for differential expression analysis [19]. Genes with a fold change greater than 2 and a corrected p-value lower than 0.05 were retained. EnhancedVolcano [20] was used to plot fold change vs p-values. Pathway enrichment analysis was performed using KEGG tool (www.genome.jp/kegg/) [14]. Heatmaps were performed in R [21] using the pheatmap package [22]. Read counts were centered and scaled for each gene to have mean zero and standard deviation one across the row.

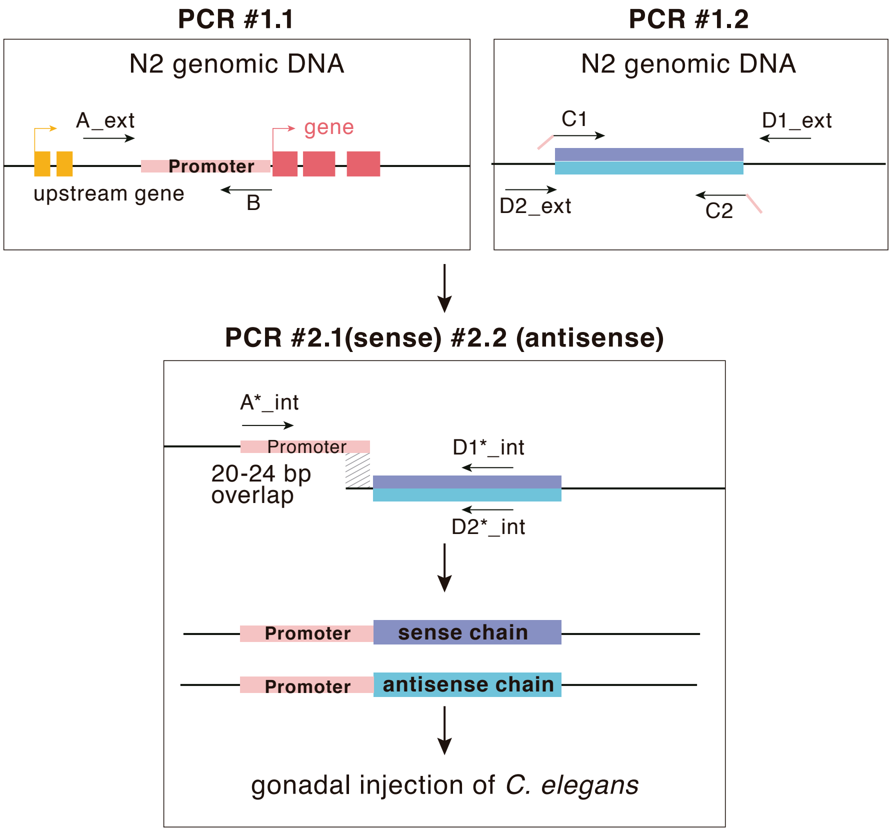


**Supplementary Figure 8. Diagram of the production of PCR-generated constructs to induce RNAi.** To assemble a promoter sequence to a gene fragment, in sense and antisense orientation, with the aim to promote tissue-specific RNAi, we used PCR fusion experiments. We used external (ext) and internal (int) primers to amplify promoter region (red rectangle), sense (purple rectangle) and antisense (blue rectangle) chain from genomic DNA. To amplify these regions, we used primers that have 20 complementary nucleotides (red tail on primers), that allow fusion by PCR using nested primers.

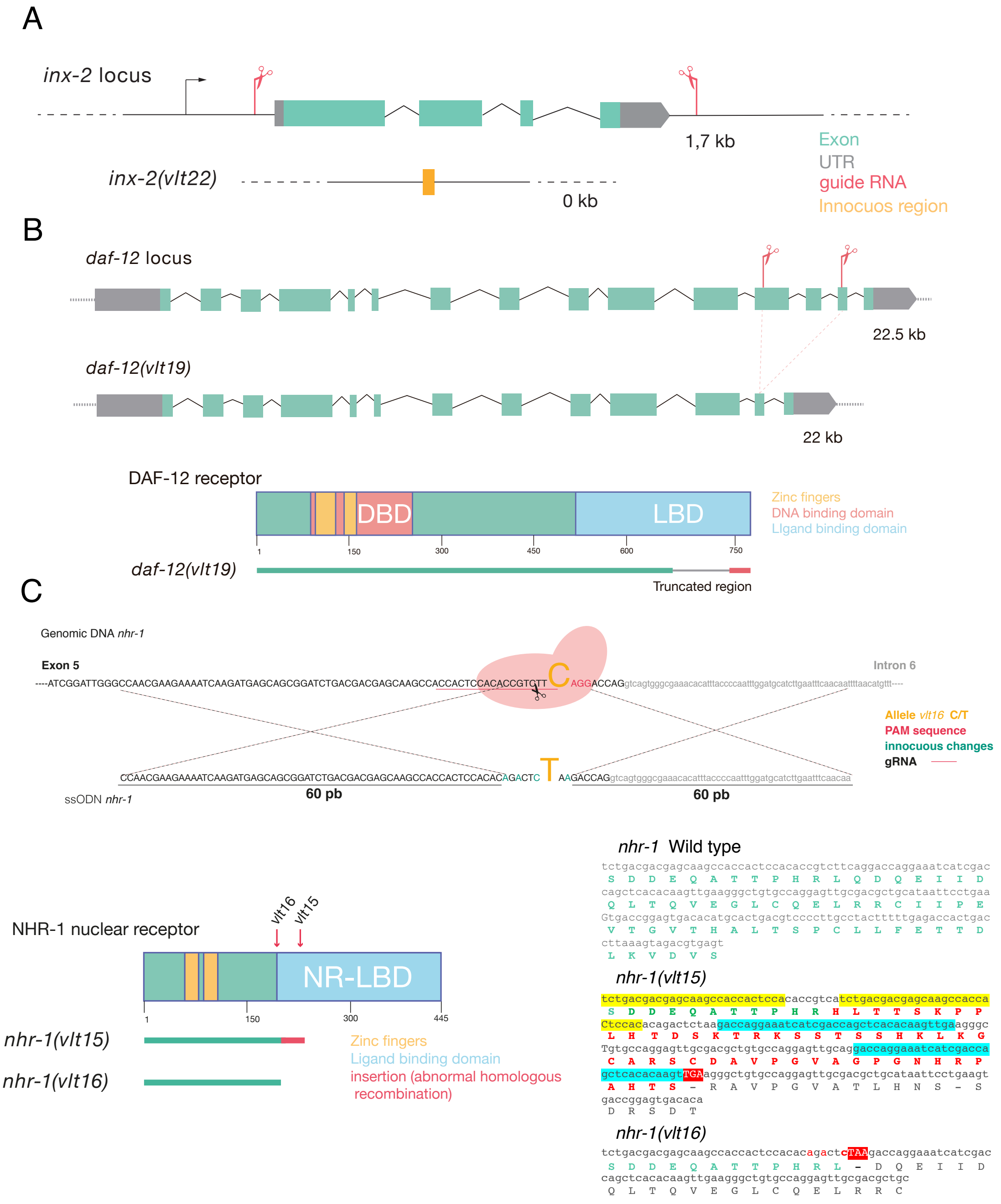


**Supplementary Figure 9. CRISPR strategies to modify gene expression.** (**A**) Disruption of the *inx-2* gene was done using two sgRNAs, that cut all coding sequences of the gene. The *vlt22* allele, obtained after the experiment, represents a complete loss of the *inx-2* gene. (**B**) *daf-12* gene disruption by the combination of two gRNAs to target exons 12 and 14. The *vlt19* deletion is a 500 bp deletion that affects in the C-terminal region of the DAF-12 protein, where it lays the ligand-binding domain (LBD) of DAF-12. (**C**) Design to introduce the *vlt16* change in *nhr-1* that emulates the allele *n6242* described by Burton *et al.*, [23]. We also obtained *vlt15* allele, resulting from an abnormal homologous recombination, which contains duplicate insertions (yellow and blue shading) that induce a frameshift (red pattern) and subsequently a TGA stop codon. The *vlt16* allele encodes a nucleotide change (C/T) that introduces a stop codon. Both alleles, *vlt16* and *vlt15*, are putative nulls.

**Supplementary Table 1.** ***C. elegans*** **strains used in this work.**

| **Strain**[1] | **Genotype** | Reference |
|---|---|---|
| Bristol N2 | *Caenorhabditis elegans* wild type background | [24] |
| AM141 | *rmIs133[unc-54p::40Q::YFP] X* | [25] |
| RVM10 | *rmIs133[unc-54p::40Q::YFP] X*; *unc-1(vlt10) X* | This work |
| CB719 | *unc-1(e719)* X | [26] |
| CB1598 | *unc-1(e1598) X* | [26] |
| RVM241 | *unc-1(vlt10) X* | This work |
| RVM23 | *rmIs133[unc-54p::40Q::YFP] X*; *unc-1(e719) X* | This work |
| RVM20 | *rmIs133[unc-54p::40Q::YFP] X*; *unc-1(e1598) X* | This work |
| AM140 | *rmIs132[unc-54p::35Q::YFP] X* | [25] |
| RVM21 | *rmIs132[unc-54p::35Q::YFP]; unc-1(e1598) X* | This work |
| RVM24 | *rmIs132[unc-54p::35Q::YFP] X; unc-1(e719) X* | This work |
| RVM26 | *rmIs132[unc-54p::35Q::YFP] X; unc-1(vlt10) X* | This work |
| NL5901 | *pkIs2386[unc-54p::α-synucleina::YFP + unc-119(+)] IV* | [27] |
| CL2006 | *dvIs2[pCL12(unc-54/Abeta human peptide 1-42 minigene)+pRF4]* | [28] |
| RVM317 | *pkIs2386[unc-54p::α-synuclein::YFP + unc-119(+)] IV; unc-1(vlt10) X* | This work |
| RVM328 | *dvIs2[pCL12(unc-54/Abeta human peptide 1-42 minigene)+pRF4], unc-1(vlt10) X* | This work |
| RVM47 | *rmIs133[unc-54p::40Q::YFP] X; unc-1(vlt10) X; vltEx47[myo-3p::unc-1(o/e); myo-2p::mCherry strain 1]* | This work |
| RVM155 | *rmIs133[unc-54p::40Q::YFP] X; unc-1(vlt10) X; vltEx155[myo-3p::unc-1(o/e); myo-2p::mCherry strain 2]* | This work |
| RVM158 | *rmIs133[unc-54p::40Q::YFP] X; unc-1(vlt10) X; vltEx158[myo-3p(o/e)); myo-2p::mCherry strain 1]* | This work |
| RVM159 | *rmIs133[unc-54p::40Q::YFP] X; unc-1(vlt10) X; vltEx159[myo-3p(o/e)); myo-2p::mCherry strain 2]* | This work |
| RVM44 | *rmIs133[unc-54p::40Q::YFP] X*; *unc-1(vlt10) X; vltEx44[rab-3p::unc-1(o/e)); myo-2p::mCherry strain 1]* | This work |
| RVM48 | *rmIs133[unc-54p::40Q::YFP] X*; *unc-1(vlt10) X; vltEx48[rab-3p::unc-1(o/e)); myo-2p::mCherry strain 2]* | This work |
| RVM42 | *rmIs133[unc-54p::40Q::YFP] X*; *vltEx42[rab-3p::AMP$^r$(RNAi)); myo-2p::mCherry strain 1]* | This work |
| RVM43 | *rmIs133[unc-54p::40Q::YFP] X; vltEx43[rab-3p:: AMP$^r$(RNAi)); myo-2p::mCherry strain 2]* | This work |
| RVM71 | *rmIs133[unc-54p::40Q::YFP] X*; *vltEx71[rab-3p::unc-1(RNAi)); myo-2p::mCherry strain 1]* | This work |
| RVM72 | *rmIs133[unc-54p::40Q::YFP] X; vltEx72[rab-3p::unc-1(RNAi)); myo-2p::mCherry strain 2]* | This work |
| RVM205 | *rmIs133[unc-54p::40Q::YFP] X; vltEx205[rab-3p(o/e)); myo-2p::mCherry strain 1]* | This work |
| RVM206 | *rmIs133[unc-54p::40Q::YFP] X; vltEx206[rab-3p(o/e)); myo-2p::mCherry strain 2]* | This work |
| RVM201 | *rmIs133[unc-54p::40Q::YFP] X; vltEx201[rab-3p::unc-1(o/e)); myo-* | This work |

| | | |
|---|---|---|
| | *2p::mCherry strain 1]* | |
| RVM221 | *rmIs133[unc-54p::40Q::YFP] X*; *vltEx221[rab-3p::unc-1(o/e)); myo-2p::mCherry strain 2]* | This work |
| RVM227 | *rmIs133[unc-54p::40Q::YFP] X*; *vltEx227[rab-3p::unc-1(n494)(o/e)); myo-2p::mCherry strain 1]* | This work |
| RVM230 | *rmIs133[unc-54p::40Q::YFP] X; vltEx230[rab-3p::unc-1(n494)(o/e)); myo-2p::mCherry strain 2]* | This work |
| RVM360 | *rmIs133[unc-54p::40Q::YFP] X; vltEx360[trx-1p::AMP*$^{r}$*(RNAi)); myo-2p::mCherry strain 1]* | This work |
| RVM361 | *rmIs133[unc-54p::40Q::YFP] X; vltEx361[trx-1p::AMP*$^{r}$*(RNAi)); myo-2p::mCherry strain 2]* | This work |
| RVM364 | *rmIs133[unc-54p::40Q::YFP] X; vltEx364[trx-1p::unc-1(RNAi)); myo-2p::mCherry strain 1]* | This work |
| RVM365 | *rmIs133[unc-54p::40Q::YFP] X; vltEx365[trx-1p::unc-1(RNAi)); myo-2p::mCherry strain 2]* | This work |
| RVM368 | *rmIs133[unc-54p::40Q::YFP] X; vltEx368[oig-1p::AMP*$^{r}$*(RNAi); osm3p::AMP*$^{r}$*(RNAi); myo-2p::mCherry strain 1]* | This work |
| RVM369 | *rmIs133[unc-54p::40Q::YFP] X; vltEx369[oig-1p::AMP*$^{r}$*(RNAi); osm-3p::AMP*$^{r}$*(RNAi); myo-2p::mCherry strain 2]* | This work |
| RVM355 | *rmIs133[unc-54p::40Q::YFP] X; vltEx355[oig-1p::unc-1(RNAi); osm-3p::unc-1(RNAi); myo-2p::mCherry strain 1]* | This work |
| RVM356 | *rmIs133[unc-54p::40Q::YFP] X; vltEx356[oig-1p::unc-1(RNAi); osm-3p::unc-1(RNAi); myo-2p::mCherry strain 2]* | This work |
| RVM362 | *rmIs133[unc-54p::40Q::YFP] X; vltEx362[glr-1p::AMP*$^{r}$*(RNAi); gpa-9p::AMP*$^{r}$*(RNAi); myo-2p::mCherry strain 1]* | This work |
| RVM363 | *rmIs133[unc-54p::40Q::YFP] X; vltEx363[glr-1p::AMP*$^{r}$*(RNAi); gpa-9p::AMP*$^{r}$*(RNAi); myo-2p::mCherry strain 2]* | This work |
| RVM366 | *rmIs133[unc-54p::40Q::YFP] X; vltEx366[glr-1p::unc-1(RNAi); gpa-9p::unc-1(RNAi); myo-2p::mCherry strain 1]* | This work |
| RVM367 | *rmIs133[unc-54p::40Q::YFP] X; vltEx367[glr-1p::unc-1(RNAi); gpa-9p::unc-1(RNAi); myo-2p::mCherry strain 2]* | This work |
| RVM386 | *rmIs133[unc-54p::40Q::YFP] X; unc-1 (vlt10) X; vltEx386[oig-1p::unc-1(o/e); myo-2p::mCherry strain 1]* | This work |
| RVM387 | *rmIs133[unc-54p::40Q::YFP] X; unc-1 (vlt10) X; vltEx387[oig-1p::unc-1(o/e); myo-2p::mCherry strain 2]* | This work |
| RVM384 | *rmIs133[unc-54p::40Q::YFP] X; unc-1 (vlt10) X; vltEx384[osm-3p::unc-1(o/e); myo-2p::mCherry strain 1]* | This work |
| RVM385 | *rmIs133[unc-54p::40Q::YFP] X; unc-1 (vlt10) X; vltEx385[osm-3p::unc-1(o/e); myo-2p::mCherry strain 2]* | This work |
| AM101 | *rmIs110[F25B3.3-p::40Q::YFP]* | [29] |
| RVM27 | *rmIs110[F25B3.3-p::40Q::YFP]; unc-1(vlt10) X* | This work |
| RVM445 | *rmIs110[F25B3.3-p::40Q::YFP]; unc-1(vlt10) X; vltEx445[osm-3p::unc-1(o/e); myo-2p::mCherry strain 1]* | This work |
| RVM446 | *rmIs110[F25B3.3-p::40Q::YFP]; unc-1(vlt10) X; vltEx445[osm-3p::unc-1(o/e); myo-2p::mCherry strain 2]* | This work |
| CB75 | *mec-2(e75) X* | [30] |
| CB138 | *unc-24(e138) IV* | [24] |
| RVM208 | *rmIs133[unc-54p::40Q::YFP] X*; *mec-2(e75) X* | This work |
| RVM192 | *rmIs133[unc-54p::40Q::YFP] X*; *unc-24(e138) IV* | This work |
| CB5 | *unc-7(e5) X* | [24] |

| | | |
|---|---|---|
| FH85 | *unc-9(ec27) X* | [31] |
| MR127 | *inx-6(rr5) IV* | [32] |
| RB1792 | *inx-7(ok2319) IV* | [33] |
| RVM209 | *rmIs133[unc-54p::40Q::YFP] X*; *unc-7(e5) X* | This work |
| RVM175 | *rmIs133[unc-54p::40Q::YFP] X*; *unc-9(ec27)* X | This work |
| RVM196 | *rmIs133[unc-54p::40Q::YFP] X*; *inx-2(vlt22) X* | This work |
| RVM197 | *rmIs133[unc-54p::40Q::YFP] X*; *inx-6(rr5) IV* | This work |
| RVM218 | *rmIs133[unc-54p::40Q::YFP] X*; *inx-7(ok2319) IV* | This work |
| RVM340 | *rmIs133[unc-54p::40Q::YFP] X; vltEx340[rab-3p::inx-6(RNAi)); myo-2p::mCherry strain 1]* | This work |
| RVM341 | *rmIs133[unc-54p::40Q::YFP] X; vltEx341[rab-3p::inx-6(RNAi)); myo-2p::mCherry strain 2]* | This work |
| RVM333 | *rmIs133[unc-54p::40Q::YFP] X; vltEx333[rab-3p::inx-2(RNAi)); myo-2p::mCherry strain 1]* | This work |
| RVM342 | *rmIs133[unc-54p::40Q::YFP] X; vltEx342[rab-3p::inx-2(RNAi)); myo-2p::mCherry strain 2]* | This work |
| RVM335 | *rmIs133[unc-54p::40Q::YFP] X; vltEx335[rab-3p::unc-7(RNAi)); myo-2p::mCherry strain 1]* | This work |
| RVM337 | *rmIs133[unc-54p::40Q::YFP] X; vltEx337[rab-3p::unc-7(RNAi)); myo-2p::mCherry strain 2]* | This work |
| RVM346 | *rmIs133[unc-54p::40Q::YFP] X; unc-7(e5) X* | This work |
| RVM337 | *rmIs133[unc-54p::40Q::YFP] X; unc-1(vlt10) X; unc-7(e5) X* | This work |
| RVM349 | *rmIs133[unc-54p::40Q::YFP] X; inx-2(vlt22) X* | This work |
| RVM348 | *rmIs133[unc-54p::40Q::YFP] X; unc-1(vlt10) X; inx-2(vlt22) X* | This work |
| CW911 | *ssu-1(fc73) V; unc-1(e580) X; rol-6(su1006)* | [6] |
| RVM142 | *rmIs133[unc-54p::40Q::YFP] X*; *ssu-1(fc73) V* | This work |
| RVM143 | *rmIs133[unc-54p::40Q::YFP] X*; *unc-1(e580)* X | This work |
| RVM144 | *rmIs133[unc-54p::40Q::YFP] X*; *ssu-1(fc73)* V; *unc-1(e580)* X | This work |
| RVM188 | *rmIs133[unc-54p::40Q::YFP] X; unc-1 (vlt10) X; vltEx188[trx-1p::AMP$^r$(RNAi)); myo-2p::mCherry strain 1]* | This work |
| RVM189 | *rmIs133[unc-54p::40Q::YFP] X; unc-1 (vlt10) X; vltEx189[trx-1p::AMP$^r$(RNAi)); myo-2p::mCherry strain 2]* | This work |
| RVM176 | *rmIs133[unc-54p::40Q::YFP] X; unc-1(vlt10) X; vltEx176[trx-1p::ssu-1(RNAi)); myo-2p::mCherry strain 1]* | This work |
| RVM232 | *rmIs133[unc-54p::40Q::YFP] X; unc-1(vlt10) X; vltEx232[trx-1p::ssu-1(RNAi)); myo-2p::mCherry strain 2]* | This work |
| VC382 | *sul-2(gk187) V* | [33] |
| RVM224 | *rmIs133[unc-54p::40Q::YFP] X*; *sul-2(gk187) V* | This work |
| RVM225 | *rmIs133[unc-54p::40Q::YFP] X; sul-2(gk187) V; unc-1(vlt10) X* | This work |
| RVM320 | *rmIs133[unc-54p::40Q::YFP] X; daf-12(vlt19) X* | This work |
| AA292 | *daf-36(k114) V* | [34] |
| RVM390 | *rmIs133[unc-54p::40Q::YFP] X; daf-36(k114) V; vltEx390[daf-36p::daf-36; myo-2p::mCherry]* | This work |
| RVM288 | *rmIs133[unc-54p::40Q::YFP] X; nhr-1(vlt15) X* | This work |
| RVM289 | *rmIs133[unc-54p::40Q::YFP] X; unc-1(vlt10) X; nhr-1(vlt15) X* | This work |

| | | |
|---|---|---|
| RVM293 | *rmIs133[unc-54p::40Q::YFP] X; nhr-1(vlt16) X* | This work |
| RVM294 | *rmIs133[unc-54p::40Q::YFP] X; unc-1(vlt10) X; nhr-1(vlt16) X* | This work |
| RVM290 | *rmIs133[unc-54p::40Q::YFP] X; unc-1(vlt10) X; nhr-1(vlt15) X; vltEx290[unc-54p::nhr-1(o/e); myo-2p::mCherry strain 1]* | This work |
| RVM291 | *rmIs133[unc-54p::40Q::YFP] X; unc-1(vlt10) X; nhr-1(vlt15) X; vltEx291[unc-54p::nhr-1(o/e); myo-2p::mCherry strain 2]* | This work |
| RVM449 | *rmIs133[unc-54p::40Q::YFP] X; unc-1(vlt10) X; nhr-1(vlt15) X; vltEx290[rab-3p::nhr-1(o/e); myo-2p::mCherry strain 1]* | This work |
| RVM450 | *rmIs133[unc-54p::40Q::YFP] X; unc-1(vlt10) X; nhr-1(vlt15) X; vltEx290[rab-3p::nhr-1(o/e); myo-2p::mCherry strain 2]* | This work |
| SJ4005 | *zcIs4[hsp-4::GFP] V* | [35] |
| RVM373 | *zcIs4[hsp-4::GFP] V; unc-1(vlt10) X* | This work |

[1]Strains are listed in order of appearance in the results.

**Supplementary Table 2. Primers used in this work.**

| PCR product | Primer sequence (5' – 3') | Strategy |
|---|---|---|
| *attB1::rab-3p ::attB5r* | Frw: **ggggacaagtttgtacaaaaaagcaggct**atcttcagatgggagcagtgg<br>Rev: **ggggacaacttttgtatacaaagttgt**catctgaaaatagggctactgtagat | Gateway MultiSite Pro |
| *attB5::unc-1 ::attB2* | Frw: **gggacaactttgtatacaaaagttga**aatgtcaaacaaggaaagaac<br>Rev: **ggggaccactttgtacaagaaagctgggta**aaggaaaatatgattattggtc | Gateway MultiSite Pro |
| *unc-1(n494)* | Frw: cttgatgaaagaactgaacat<br>Rev: atgttcagttctttcatcaag | Gateway MultiSite Pro |
| *rab-3p* | A: M13F<br>B: catctgaaaatagggctactgtagat | Tissue-specific RNAi PCR #1.1 |
| *unc-1 _sense/antisense* overlapping: *rab-3p* | C1: **atctacagtagccctattttcagatg**atgtcaaacaaggaaagaacagag<br>D1: M13R<br>C2: **atctacagtagccctattttcagatg**ttattggtctttttcataaatgctcc<br>D2: M13F<br>A*: atcttcagatgggagcagtgg<br>D1*: aaggaaaatatgattattggtc<br>D2*: aaatgtcaaacaaggaaagaac | Tissue-specific RNAi<br>PCR #1.2<br>*PCR #2.1 |
| *AMP^r _sense/antisense* overlapping: *rab-3p* | C1: **atctacagtagccctattttcagatg**gcatcttacggatggcatgacag<br>D1: acagagttcttgaagtggtggc<br>C2: **atctacagtagccctattttcagatg**cgtttggtatggcttcattcagc<br>D2: gcttacagacaagctgtgaccg<br>A*: atcttcagatgggagcagtgg<br>D1*: cgtttggtatggcttcattcagc<br>D2*: gcatcttacggatggcatgacag | Tissue-specific RNAi<br>PCR #1.2<br>*PCR #2.1 |
| *inx-2 _sense/antisense* overlapping: *rab-3p* | C1: **atctacagtagccctattttcagatg**gtaccacttgtccttcttttccaag<br>D1: gtcaaggaaatgtttcagaagaacc<br>C2: **atctacagtagccctattttcagatg**gtcaaggaaatgtttcagaagaacc<br>D2: gtaccacttgtccttcttttccaag<br>A*: atcttcagatgggagcagtgg<br>D1*: atttgcagaatgcaattgacaa<br>D2*: ctgccatgtttgtgctcccctta | Tissue-specific RNAi<br>PCR #1.2<br>*PCR #2.1 |
| *inx-6 _sense/antisense* overlapping: *rab-3p* | C1: **atctacagtagccctattttcagatg**gtcaactttgtgaaccagtact<br>D1: aatgtatacagatggacgtcct<br>C2: **atctacagtagccctattttcagatg**aatgtatacagatggacgtcct<br>D2: gtcaactttgtgaaccagtact<br>A*: atcttcagatgggagcagtgg | Tissue-specific RNAi<br>PCR #1.2<br>*PCR #2.1 |

| | | |
|---|---|---|
| | D1*: tcgaaagctgctagccgagttttg<br>D2*: tgttccactcgaccagcaactagc | |
| *unc-7 _sense/antisense* overlapping: *rab-3p* | C1: **atctacagtagccctattttcagatg**ctttgagcactcaaagaaaaactct<br>D1: tcagtctatcgtcccttgaccgtgt<br>C2: **atctacagtagccctattttcagatg**tcagtctatcgtcccttgaccgtgt<br>D2: ctttgagcactcaaagaaaaactct<br>A*: atcttcagatgggagcagtgg<br>D1*: tccgcatcccaaaattgcgtcgg<br>D2*: actataattcaaaagcaacctaatg | Tissue-specific RNAi<br>PCR #1.2<br>*PCR #2.1 |
| *trx-1p* | A: aaccaattgagttggcacttcg<br>B: aaccttggtgagagacatgatg | Tissue-specific RNAi PCR #1.1 |
| *ssu-1 _sense/antisense* overlapping: *trx-1p* | C1: **catcatgtctctcaccaaggtt**ccagagctctgtgtgcaatcgc<br>D1: catgtttcgcgtatttttctgc<br>C2: **catcatgtctctcaccaaggtt**ccttatacacagcattttcc<br>D2: cgtggcgggacccaaaatctc<br>A*: agaatggatacctgatcatt<br>D1*: ccttatacacagcattttcc<br>D2*: ccagagctctgtgtgcaatcgc | Tissue-specific RNAi<br>PCR #1.2<br>*PCR #2.1 |
| *$AMP^r$ _sense/antisense* overlapping: *trx-1p* | C1: **catcatgtctctcaccaaggtt**gcatcttacggatggcatgacag<br>D1: acagagttcttgaagtggtggc<br>C2: **catcatgtctctcaccaaggtt**cgtttggtatggcttcattcagc<br>D2: gcttacagacaagctgtgaccg<br>A*: agaatggatacctgatcatt<br>D1*: cgtttggtatggcttcattcagc<br>D2*: gcatcttacggatggcatgacag | Tissue-specific RNAi<br>PCR #1.2<br>*PCR #2.1 |
| *unc-1 _sense/antisense* overlapping: *trx-1p* | C1: **catcatgtctctcaccaaggtt**atgtcaaacaaggaaagaacagag<br>D1: M13R<br>C2: **catcatgtctctcaccaaggtt**ttattggtctttttcataaatgctcc<br>D2: M13F<br>A*: agaatggatacctgatcatt<br>D1*: aaggaaaatatgattattggtc<br>D2*: aaatgtcaaacaaggaaagaac | Tissue-specific RNAi<br>PCR #1.2<br>*PCR #2.1 |
| *oig-1p* | A: agagcaagcagtcagtgaaaatgt<br>B: agtcgaacgttttgcagaattatg | Tissue-specific RNAi PCR #1.1 |
| *osm-3p* | A: gcttaaaatccggctaaaattca<br>B: tccgacgcatagctggaaattttg | Tissue-specific RNAi PCR #1.1 |
| *glr-1p* | A: aacaagaaagtcgtagttgttac<br>B: tgtgaatgtgtcagattgggtgcc | Tissue-specific RNAi PCR #1.1 |
| *gpa-9p* | A: gatggtcccggaaaacatcatcg<br>B: cccattgcatatttcattaaac | Tissue-specific RNAi PCR #1.1 |
| *unc-1 _sense* overlapping: *oig-1p* | C1: **cataattctgcaaaacgttcgact**atgtcaaacaaggaaagaacagag<br>D1: M13R<br>A*: aacatgttttggagcatatttcgcg<br>D1*: aaggaaaatatgattattggtc | Tissue-specific RNAi<br>PCR #1.2<br>*PCR #2.1 |
| *unc-1 _antisense* overlapping: *osm-3p* | C2: **caaaatttccagctatgcgtcgga**ttattggtctttttcataaatgctcc<br>D2: M13F<br>A*: aattaaatttgcctgaaaatccg<br>D2*: aaatgtcaaacaaggaaagaac | Tissue-specific RNAi<br>PCR #1.2<br>*PCR #2.1 |
| *unc-1 _sense* overlapping: *glr-1p* | C1:**ggcacccaatctgacacattcaca**atgtcaaacaaggaaagaacagag<br>D1: M13R<br>A*: aataattataagagacgtgtag<br>D1*: aaggaaaatatgattattggtc | Tissue-specific RNAi<br>PCR #1.2<br>*PCR #2.1 |
| *unc-1 _antisense* overlapping: *gpa-9p* | C2: **gtttaatgaaatatgcaatgggtt**attggtctttttcataaatgctcc<br>D2: M13F<br>A*: accgaatcaaaatatctgaat<br>D2*: aaatgtcaaacaaggaaagaac | Tissue-specific RNAi<br>PCR #1.2<br>*PCR #2.1 |
| *$AMP^r$ _sense* overlapping: *oig-1p* | C1: **cataattctgcaaaacgttcgact**gcatcttacggatggcatgacag<br>D1: acagagttcttgaagtggtggc<br>A*: aacatgttttggagcatatttcgcg<br>D1*: cgtttggtatggcttcattcagc | Tissue-specific RNAi<br>PCR #1.2<br>*PCR #2.1 |

| | | |
|---|---|---|
| *AMP^r _antisense* overlapping: *osm-3p* | C2: **caaaatttccagctatgcgtcgga**cgtttggtatggcttcattcagc<br>D2: gcttacagacaagctgtgaccg<br>A*: aattaaatttgcctgaaaatccg<br>D2*: gcatcttacggatggcatgacag | Tissue-specific RNAi<br>PCR #1.2<br>*PCR #2.1 |
| *AMP^r _sense* overlapping: *glr-1p* | C1: **ggcacccaatctgacacattcaca**gcatcttacggatggcatgacag<br>D1: acagagttcttgaagtggtggc<br>A*: aataattataagagacgtgtag<br>D1*: cgtttggtatggcttcattcagc | Tissue-specific RNAi<br>PCR #1.2<br>*PCR #2.1 |
| *AMP^r _antisense* overlapping: *gpa-9p* | C2: **gtttaatgaaatatgcaatggg**cgtttggtatggcttcattcagc<br>D2: gcttacagacaagctgtgaccg<br>A*: accgaatcaaaatatctgaat<br>D2*: gcatcttacggatggcatgacag | Tissue-specific RNAi<br>PCR #1.2<br>*PCR #2.1 |
| *unc-1(cDNA) ::unc-54t* overlapping: *oig-1p* | C1:**cataattctgcaaaacgttcgact**atgtcaaacaaggaaagaacagag<br>D1: aaacagttatgtttggtatattg<br>A*: aacatgttttggagcatatttcgcg<br>D1*: aatgtattctgtcatttaaggc | Tissue-specific Rescue<br>PCR #1.2<br>*PCR #2.1 |
| *unc-1(cDNA) ::unc-54t* overlapping: *osm-3p* | C1:**caaaatttccagctatgcgtcgga**atgtcaaacaaggaaagaacagag<br>D1: aaacagttatgtttggtatattg<br>A*: aattaaatttgcctgaaaatccg<br>D1*: aatgtattctgtcatttaaggc | Tissue-specific Rescue<br>PCR #1.2<br>*PCR #2.1 |
| *sul-1* | Frw: ttcagctctggaaatgctactac<br>Rev: tctttttgtacttcaagtagtgc | Feeding RNAi |
| *sul-2* | Frw: gctcacctagcagagctggattc<br>Rev: gtctctcgtgttgatcttgtgcc | Feeding RNAi |
| *sul-3* | Frw: ttacatacaccaaacctccggc<br>Rev: ttgaagtaacccgtttgtggtcc | Feeding RNAi |
| *elo-2* | Frw: tcatgcactggtatcatcatgcc<br>Rev: gcgcaaccaggaacagaatcag | Feeding RNAi |
| *fat-6* | Frw: cgtaagcatccacaagttaagg<br>Rev: ggtaattgaggaatcgtatggc | Feeding RNAi |
| *skn-1* | Frw: gatcgcgagagttgttccactgg<br>Rev: ggctttaataaggtttcgaccgag | Feeding RNAi |
